# Human iPSC-Based Model Reveals NOX4 as Therapeutic Target in Duchenne Cardiomyopathy

**DOI:** 10.1101/2021.09.13.460090

**Authors:** Robin Duelen, Domiziana Costamagna, Guillaume Gilbert, Liesbeth De Waele, Nathalie Goemans, Kaat Desloovere, Catherine M. Verfaillie, Karin R. Sipido, Gunnar M. Buyse, Maurilio Sampaolesi

## Abstract

Duchenne muscular dystrophy (DMD) is an X-linked progressive muscle disorder, caused by mutations in the *Dystrophin* gene. Cardiomyopathy is one of the major causes of early death. In this study, we used DMD patient-specific induced pluripotent stem cells (iPSCs) to model cardiomyopathic features in DMD and unravel novel pathological mechanistic insights. Cardiomyocytes (CMs) differentiated from DMD iPSCs showed enhanced premature cell death, due to significantly elevated intracellular reactive oxygen species (ROS) concentrations, as a result of depolarized mitochondria and high NADPH oxidase 4 (NOX4) protein levels. Genetic correction of *Dystrophin* through CRISPR/Cas9 editing restored normal ROS levels. Application of ROS reduction by N-acetyl-L-cysteine (NAC), partial Dystrophin re-expression by ataluren (PTC124) and enhancing mitochondrial electron transport chain function by idebenone improved cell survival of DMD iPSC-CMs. We show applications that could counteract the detrimental oxidative stress environment in DMD iPSC-CMs by stimulating adenosine triphosphate (ATP) production. ATP could bind to the ATP-binding domain in the NOX4 enzyme, and we demonstrate that ATP resulted in partial inhibition of the NADPH-dependent ROS production of NOX4.

Considering the complexity and the early cellular stress responses in DMD cardiomyopathy, we propose to target ROS production and prevent the detrimental effects of NOX4 on DMD CMs as a promising therapeutic strategy.

**GRAPHICAL ABSTRACT:** 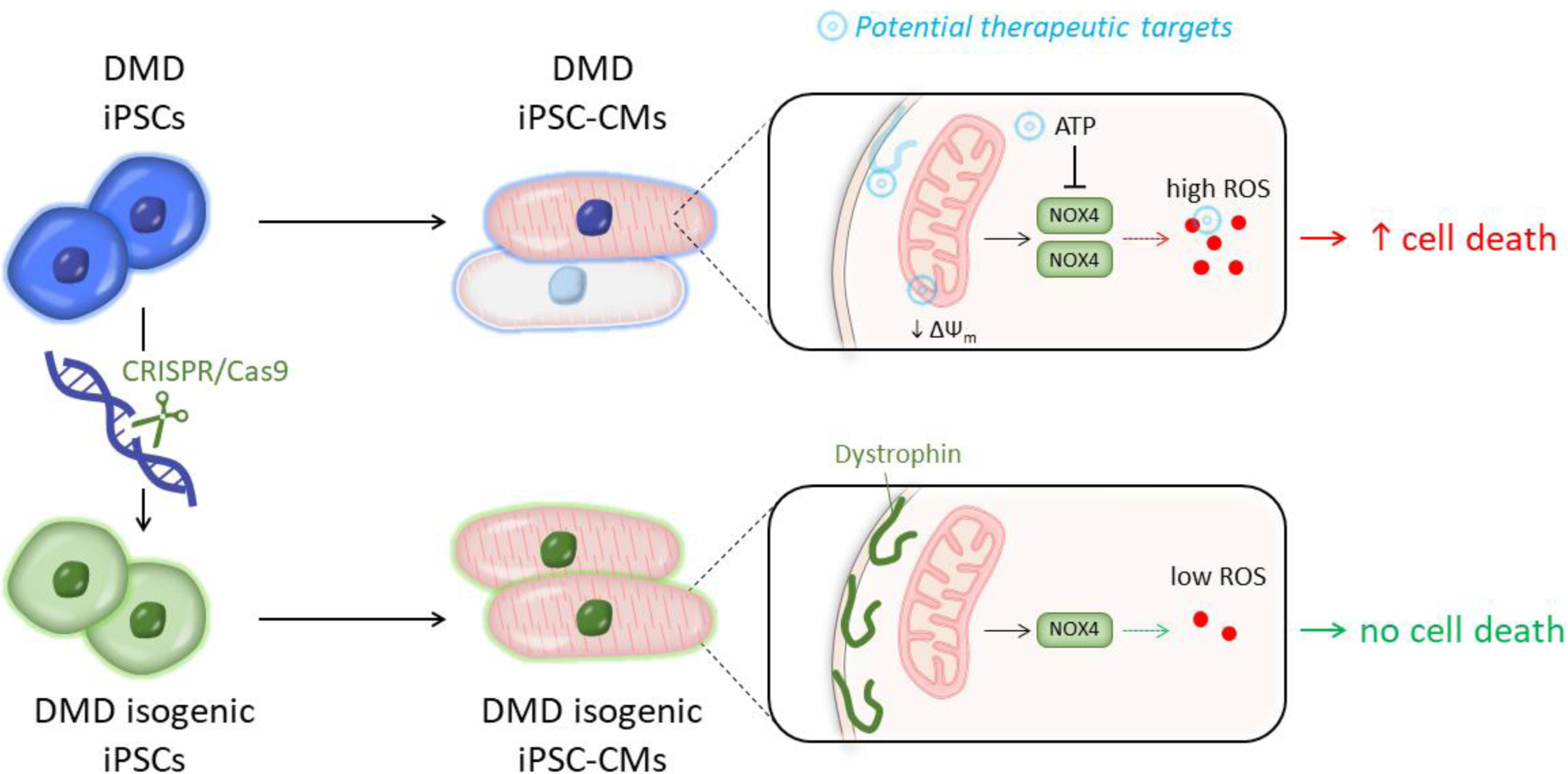
The use of human induced pluripotent stem cell-derived cardiomyocytes (iPSC-CMs) from Duchenne muscular dystrophy (DMD) patients to model cardiomyopathic features in DMD and unravel novel pathological mechanistic insights. DMD iPSC-CMs showed accelerated cell death, caused by increased intracellular reactive oxygen species (ROS) levels. By intervention at different target sites, beneficial effects on the mitochondrial membrane potential (ΔΨm) and the expression and ROS-producing activity of the cardiac-specific NADPH-oxidase 4 isoform (NOX4) were observed, resulting in an increased cell survival and function of DMD iPSC-CMs.

## INTRODUCTION

The shortage of human cardiac cell sources has challenged cardiovascular disease modeling and drug development. The generation of functional cardiomyocytes (CMs) differentiated from human induced pluripotent stem cells (iPSCs) overcomes current limitations and offers an extraordinary platform to develop iPSC-based models to study the genetic disease phenotype of cardiomyopathic pathologies *in vitro* (*1, 2*).

Mutations in the *Dystrophin* gene cause the X-linked disorder Duchenne muscular dystrophy (DMD), the most common and severe phenotype among the muscular dystrophies (*3*). Most DMD patients develop adverse myocardial remodeling and chronic cardiomyopathy, a major cause of morbidity and early mortality (*4*). With current standards of care, the median life expectancy at birth in DMD seems to have improved considerably during the last decades, ranged between 21.0 and 39.6 years (*5*).

The Dystrophin protein has a crucial role during muscle contraction and stretch. Loss of function or absence lead to myocyte sarcolemma instability during contraction-relaxation cycles, making myocytes more susceptible to stretch-induced damage and necrosis (*6*). The signaling-mediated role of Dystrophin and the associated dystrophin glycoprotein complex is not yet fully understood (*7*). The pathophysiological role of Dystrophin in the heart is poorly defined and little is known about the earliest stages of DMD cardiomyopathy. Multiple pathways are involved including dysregulation of calcium (Ca^2+^) homeostasis, oxidative stress, inflammation and functional ischemia.

Oxidative stress is involved in the pathogenesis of heart failure. However, clinical trials using antioxidants have shown limited success (*8, 9*). Nicotinamide adenine dinucleotide phosphate (NADPH) oxidase (NOX) family enzymes generate reactive oxygen species (ROS) in a highly regulated manner, modulating several physiological aspects such as host defense, posttranslational processing of proteins, cellular signaling, regulation of gene expression and cell differentiation (*10, 11*). However, NOX family enzymes also contribute to a wide range of pathological processes, in particular cardiovascular diseases (*12-16*). The NOX4 isoform is predominantly expressed in CMs, although the precise location remains controversial. It is constitutively active at low level (*17*), inducing cardioprotective effects under chronic stress (*18*). The exact role of NOX4 in CMs is still not clear, even though high levels of NOX4 could have severe detrimental effects (*12-16*). Targeting NOX isoforms may be a useful therapeutic strategy. Therefore, there has been significant focus on the potential role of ROS-generating NOX isoform proteins in the pathogenesis of DMD (*14, 15, 19*).

Several innovative therapeutic approaches focus on targeting the primary defect such as restoring the function or expression of Dystrophin through exon skipping (*20*), ribosomal read-through technology (*21*), as well as gene (*22*) and cell therapy (*23*). Recent technological breakthroughs in genome editing successfully enabled the correction of the genetic mutation (*24, 25*). In addition, compounds targeting downstream pathophysiology are under investigation in clinical trials (*26*). Idebenone (2,3-dimethoxy-5-methyl-6-(10-hydroxy)decyl-1,4-benzoquinone), a synthetic analogue of coenzyme Q10, has a dual mode-of-action. First, it detoxifies ROS by donating electrons to produce non-toxic reaction products. Second, it donates electrons directly to Complex III of the mitochondrial electron transport chain (ETC), which restores electron flow, proton pumping activity of Complexes III and IV, and adenosine triphosphate (ATP) production by Complex V. Phase 2 and phase 3 randomized placebo-controlled trials have demonstrated a beneficial role of idebenone in DMD patients (*27, 28*).

In this study, we used DMD patient-specific iPSC-derived CMs (iPSC-CMs) to model cardiomyopathic features in DMD and to explore pathological mechanisms. We observed mitochondrial dysfunction and increased concentrations of intracellular ROS in DMD iPSC-CMs, due to significantly increased expression and activity of NOX4. These features were not present in CRISPR/Cas9 genetically corrected DMD isogenic iPSC-CMs. Additionally, by administration of the ROS scavenger N-acetyl-L-cysteine (NAC), the read-through chemical drug ataluren (PTC124) or the synthetic benzoquinone idebenone to differentiated DMD iPSC-CMs, we observed beneficial outcomes regarding cell survival and function. Interestingly, idebenone showed superior improvements compared to NAC and PTC124 alone or in combination. In addition, our data indicated that idebenone counteracted the hyperactive ROS-producing activity of the cardiac-specific NOX4 isoform through a mechanism mediated by ATP production that could reduce NOX4 activity, resulting in an improved contractile function.

In conclusion, using DMD patient-derived iPSCs, we established an *in vitro* model to recapitulate DMD heart disease phenotypes and to study novel molecular disease mechanisms that might become interesting therapeutic targets for cardiomyopathy in DMD patients.

## RESULTS

### Generation of Integration-Free DMD iPSCs

To obtain an unlimited cell source of CMs, recapitulating aspects of a single-gene disease phenotype, iPSC lines were generated from human dermal fibroblasts (hFs) and human peripheral blood mononuclear cells (hPBMCs) obtained from DMD patients with known *Dystrophin* mutations (Table S1). Somatic cells were reprogrammed towards a pluripotent state using the integration-free Sendai virus (SeV) vectors (Fig. S1A-C), which expressed the OSKM (OCT3/4, SOX2, KLF4 and c-MYC) pluripotency markers. Subcutaneously injected DMD iPSC lines into immunodeficient mice displayed teratoma formation, successfully showing the differentiation capacity into all three developmental germ layers (ectoderm, mesoderm and endoderm; Fig. S1D-E). Furthermore, detailed pluripotency analysis for related genes and proteins is given in Supplemental Information (Fig. S2A-B). Three human control iPSC lines were used, generated from healthy donors with no neuromuscular disorders (Table S1).

### CRISPR/Cas9-Mediated Correction of Nonsense Mutation in Dystrophin Gene

Additionally, we created an isogenic control line to exclude genetic background variability. The isogenic control line was generated using CRISPR/Cas9 technology from the DMD iPSC patient line, characterized by a genetic point mutation in exon 35 (c.4,996C > T; p.Arg1,666X) of the *Dystrophin* gene, resulting in a premature stop codon and consequently in the complete absence of a functional Dystrophin protein (Fig. 1A). To restore full-length expression of the *Dystrophin* gene, two 20-nucleotide single-guide RNAs (sgRNAs) were designed to induce Cas9-mediated double-stranded breaks (DSB) in the genomic DNA of the Dystrophin-deficient iPSCs (Fig. 1B). SgRNA specificity and CRISPR/Cas9 DSB cutting were evaluated in HEK293T cells by the appearance of non-homologous end joining (NHEJ) events after transfection of the sgRNA-Cas9 plasmids (Fig. S3A-B). Cas9-mediated genome editing was performed via homology-directed repair (HDR), using a plasmid-based donor repair template with homology arm regions for the *Dystrophin* gene exon of interest, in order to substitute the premature stop codon into the original amino acid codon for arginine (Fig. 1B). Sequencing analysis of exon 35 of the *Dystrophin* gene confirmed CRISPR/Cas9-mediated genomic correction, further indicated as DMD isogenic control (Fig. 1C). CRISPR/Cas9 off-target events were analyzed based on sequence homology of sgRNAs (Fig. S3C) and detailed comparative genomic hybridization (CGH) molecular karyotyping did not show additional chromosomal abnormalities due to unwanted Cas9-mediated DSB cuts (Fig. S3D). To demonstrate that gene editing did not influence the pluripotency state of the DMD isogenic control, pluripotency genes (*c-MYC, GDF-3, KLF4, NANOG, OCT4, REX1, SOX2* and *hTERT*) and proteins (OCT4, NANOG, SSEA4, SOX2, TRA-1-60 and LIN28) were analyzed in several undifferentiated human pluripotent stem cell (PSC) lines (Fig. S2A-B). Furthermore, immunofluorescent staining showed the expression of Dystrophin protein levels (green) in differentiated DMD iPSC-CMs (cTnT, red and Hoechst, blue) after CRISPR/Cas9 correction (Fig. 1D).

**Fig. 1:**
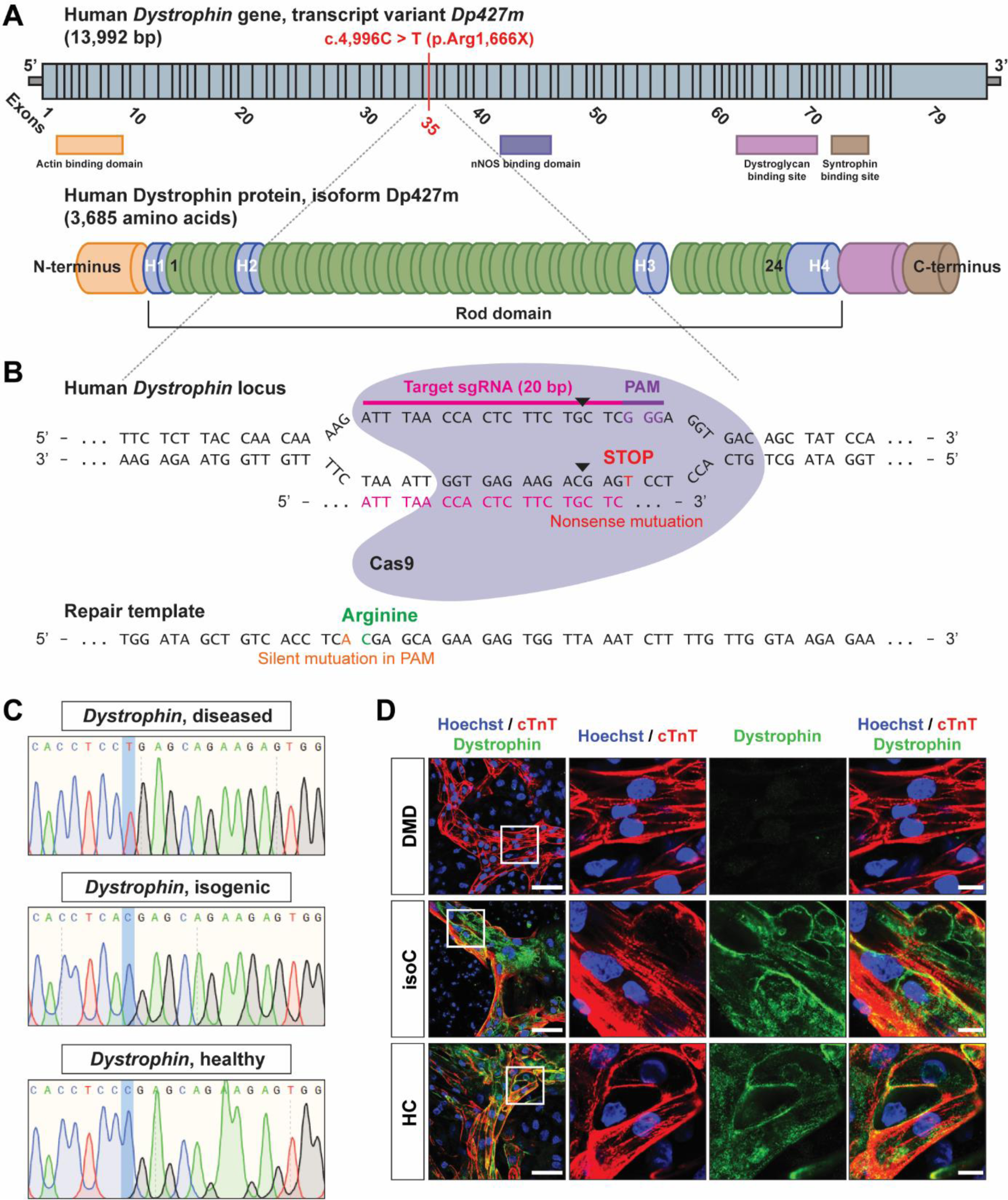
DMD iPSC CRISPR/Cas9 gene editing of a nonsense mutation in exon 35 (c.4,996C > T; p.Arg1,666X) of the *Dystrophin* gene. **(A)** Schematic representation of the human *Dystrophin* gene sequence (*top,* transcript variant *Dp427m*) and the encoded Dystrophin protein (*bottom,* isoform Dp427m). The genetic point mutation is located in exon 35 of the *Dystrophin* gene, resulting in a premature stop codon. **(B)** The 20-nucleotide sgRNA (ATTTAACCACTCTTCTGCTC) to induce the Cas9-mediated DSB (indicated as black triangles). The donor repair template, containing the CRISPR/Cas9-mediated genetic correction of the nonsense mutation in the *Dystrophin* gene. **(C)** DNA sequencing of the mutated region of interest of the *Dystrophin* gene before (DMD diseased) and after (DMD isogenic) CRISPR/Cas9 gene editing. **(D)** Immunofluorescent staining showing the expression of Dystrophin protein levels (green) in differentiated DMD iPSC-CMs (cTnT, red and Hoechst, blue) after CRISPR/Cas9-mediated genetic correction. Scale bar: 50 μm. White boxes with corresponding insets at higher magnification. Scale bar: 10 μm.

### Human iPSC-CMs to Model Diseased Heart Phenotype in DMD

Burridge *et al.* developed a fully chemically defined and small molecule based cardiac differentiation protocol, effective for several human iPSC lines and with high yield of mainly ventricular-like CMs (*29*). Here, we differentiated control and DMD iPSC lines to CMs, according to this monolayer-based cardiac differentiation strategy (*29*), with additional 3D maturation in fibrin-based engineered heart tissue (EHT) constructs (Fig. 2A-B) (*30*). During the early phases of cardiac differentiation, human iPSCs were treated with chemical Wnt signaling mediators (CHIR99021 and IWR-1) to obtain high CM yields (Fig. 2A). Additional 3D maturation of iPSC-CMs could significantly increase the expression of the cardiac-specific maturation isoforms *MYL2* and *TNNI3* (Fig. 2C). Immunostaining of cTnT positive iPSC-CMs additionally matured in 3D EHTs showed structural aligned orientation due to mechanical loading of the flexible microposts compared to classical 2D monolayer-based differentiation systems (Fig. 2D). Importantly, differentiated iPSC-CMs from DMD patients manifested pathologic features of cardiac involvement. They exhibited a significant reduction of the L-type Ca^2+^ current, indicating abnormal Ca^2+^ homeostasis (Fig. 2E), and representative action potential (AP) recordings from DMD iPSC-CMs displayed arrhythmogenic firing pattern including delayed afterdepolarizations (DADs) and oscillatory prepotentials (OPPs; Fig. 2F), as reported in literature by Eisen *et al.* and others (*31-36*). Furthermore, detailed patch-clamp recordings at day 24 of differentiation showed significant longer mean action potential duration at 90% repolarization (APD90) of DMD iPSC-CMs compared to control iPSC-CMs (Fig. 2G). Other electrophysiological parameters including AP amplitude, resting membrane potential (RMP), cell capacitance and beating frequency did not show significant differences (Fig. 2H).

**Fig. 2:**
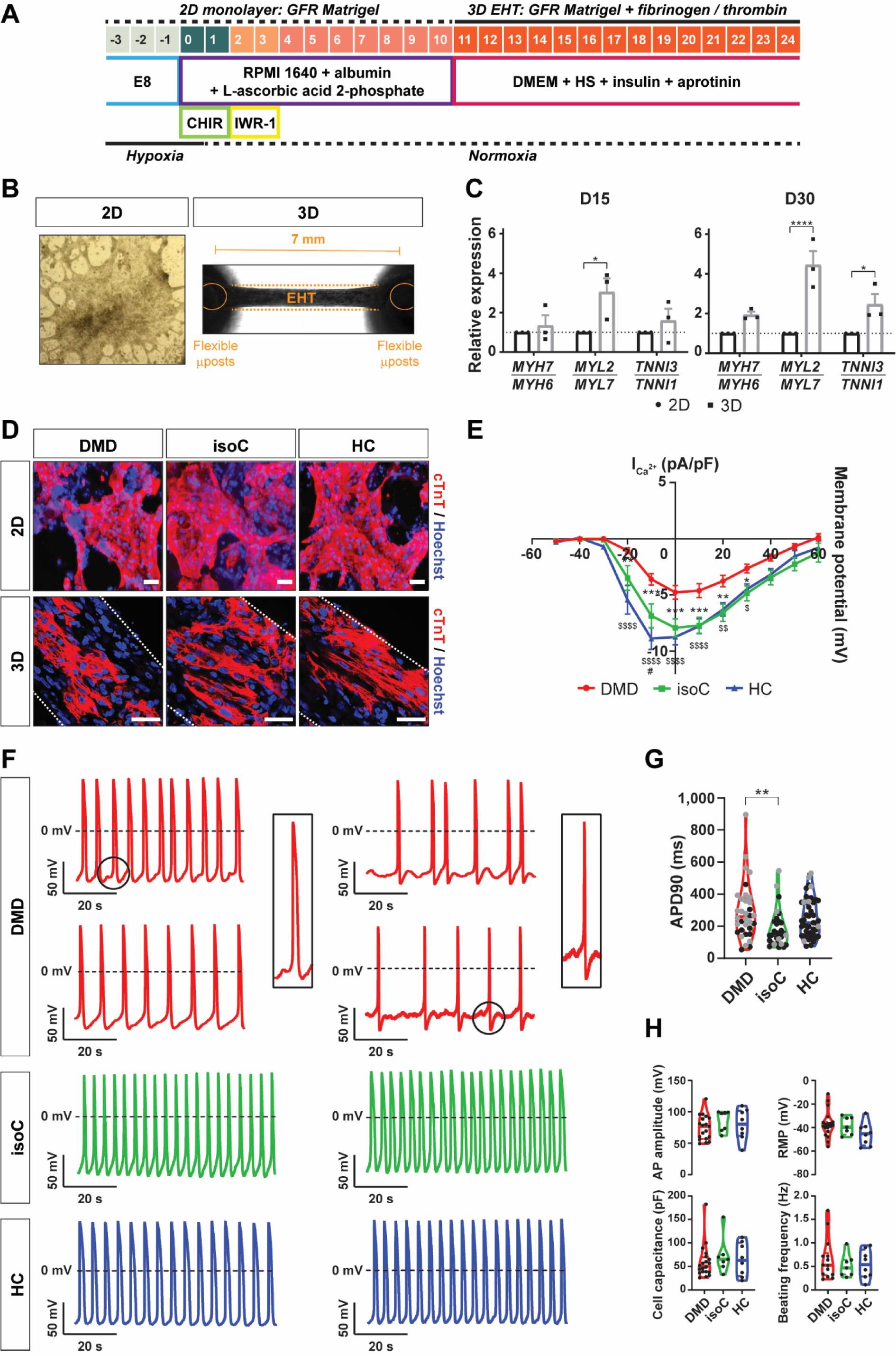
Characterization of the iPSC-CM differentiation protocol. **(A)** Schematic representation of the cardiac differentiation protocol. Human iPSCs were differentiated to CMs in a monolayer cardiac differentiation protocol, using chemical Wnt signaling mediators (CHIR99021 and IWR-1), and, eventually, further matured into 3D EHT constructs based on fibrinogen and thrombin polymerization. **(B)** Representative example of 2D monolayer-based cardiac differentiation (*left panel*) and 3D mini-EHT construct between two flexible microposts, positioned 7 mm from each other (*right panel*). **(C)** Normalized gene expression ratios for isoforms of *Myosin Heavy Chain* (*MYH7/MYH6*), *Myosin Light Chain* (*MYL2/MYL7*) and *Cardiac Troponin I* (*TNNI3/TNNI1*) after 15 and 30 days of differentiation. Data were representative of three independent experiments (N = 3) and values were expressed as mean ± SEM. Significance of the difference was indicated as follows: *P < 0.05; **P < 0.01; ***P < 0.001 and ****P < 0.0001. **(D)** Immunostaining of Cardiac Troponin C (cTnT) positive CMs (cTnT, red and Hoechst, blue) in monolayer-based cardiac differentiation (2D) or EHT constructs (3D). White dotted lines indicated the borders of the 3D EHT constructs. Scale bar: 50 μm. **(E)** Voltage-current relation curve of the L-type Ca^2+^ current (pA/pF), assessed after whole-cell patch-clamp configuration. Data were representative of three independent experiments (N = 3; DMD: n = 13, DMD isogenic: n = 7, healthy: n = 11) and values were expressed as mean ± SEM. Significance of the difference was indicated as follows: *P < 0.05; **P < 0.01; ***P < 0.001 and ****P < 0.0001 (DMD vs. DMD isogenic control); or ^$^P < 0.05; ^$$^P < 0.01; ^$$$^P < 0.001 and ^$$$$^P < 0.0001 (DMD vs. healthy control). **(F)** Representative AP recordings from DMD and control iPSC-CMs. DMD iPSC-CMs displayed arrhythmogenic firing pattern including DADs and OPPs. **(G)** Patch-clamp recordings at day 24 of differentiation for mean APD90 (ms). Additional measurements were performed with di-4-ANEPPS (gray dots). **(H)** Patch-clamp recordings for AP amplitude (mV), RMP (mV), cell capacitance (pF) and beating frequency (Hz). Data were representative of three independent experiments (N = 3; DMD: n = 19, DMD isogenic: n = 7, healthy: n = 8). Significance of the difference was indicated as follows: *P < 0.05; **P < 0.01; ***P < 0.001 and ****P < 0.0001.

### Enhanced Cell Death and Excessive Intracellular ROS Levels in DMD iPSC-CM Cultures

The absence of Dystrophin protein in differentiated iPSC-CMs from DMD patients (Fig. S4) results in progressive loss of CMs (*6, 32*). In this study, we wanted to identify novel pathological cues that caused decreased cell survival of DMD iPSC-CMs. We mainly used the DMD iPSC patient line that was characterized by the nonsense mutation in exon 35 (c.4,996C > T; p.Arg1,666X) of the *Dystrophin* gene (DMD #2 in Table S1). This DMD line represents a subgroup of DMD patients (approximately 13%) that is responsive to the read-through chemical drug ataluren (PTC124; Fig. 5A-C). Cell death was examined by flow cytometric analyses, using annexin V and 7-amino-actinomycin D (7AAD). DMD iPSC-CMs underwent accelerated cell death (DMD, early apoptosis: 15 ± 1% and late apoptosis: 33 ± 6%) compared to corresponding DMD isogenic (isoC, respectively 3 ± 0% and 7 ± 1%) and healthy controls (HC, respectively 5 ± 0% and 15 ± 1%; Fig. 3A and Fig. S6A, *left panels*). A remarkable percentage of DMD iPSC-CMs had high intracellular ROS concentrations (DMD, [ROS]^high^: 68 ± 2%) compared to controls (isoC, [ROS]^high^: 46 ± 3% and HC, [ROS]^high^: 41 ± 1%; Fig. 3B and Fig. S6A, *middle panels*). Moreover, the intracellular ROS content (mean fluorescence intensity, MFI) in DMD iPSC-CMs was significantly higher (DMD, 28,924 ± 1,864 vs. isoC, 5,276 ± 254 and vs. HC, 6,198 ± 213; Fig. 3C and Fig. S6A, *right panels*). Upon treatment with NAC and PTC124 (alone or in combination) as well with idebenone, DMD iPSC-CMs showed increased cell survival (DMD early apoptosis, NAC: 7 ± 1%, PTC124: 8 ± 1%, NAC+PTC124: 9 ± 1% and idebenone: 3 ± 0% vs. untreated: 15 ± 1% and DMD late apoptosis, NAC: 25 ± 2%, PTC124: 24 ± 2%, NAC+PTC124: 21 ± 2% and idebenone: 5 ± 1% vs. untreated: 33 ± 6%; Fig. 3A and Fig. S6A, *left panels*) and reduced intracellular ROS levels (DMD [ROS]^high^, NAC: 36 ± 4%, PTC124: 41 ± 11%, NAC+PTC124: 48 ± 7% and idebenone: 52 ± 3% vs. untreated: 68 ± 2%; Fig. 3B and Fig. S6A, *middle panels*) compared to untreated DMD iPSC-CMs. The specificity of the drug effect on the CM death and on the intracellular ROS levels of the experimental groups is shown in Supplemental Information (Fig. S7A-D). Taken together, these results show increased intracellular ROS levels in DMD iPSC-CMs. Interestingly, NAC, PTC124 and idebenone had beneficial effects on the cell survival, although idebenone addition exhibited superior effects on DMD iPSC-CM cultures.

**Fig. 3:**
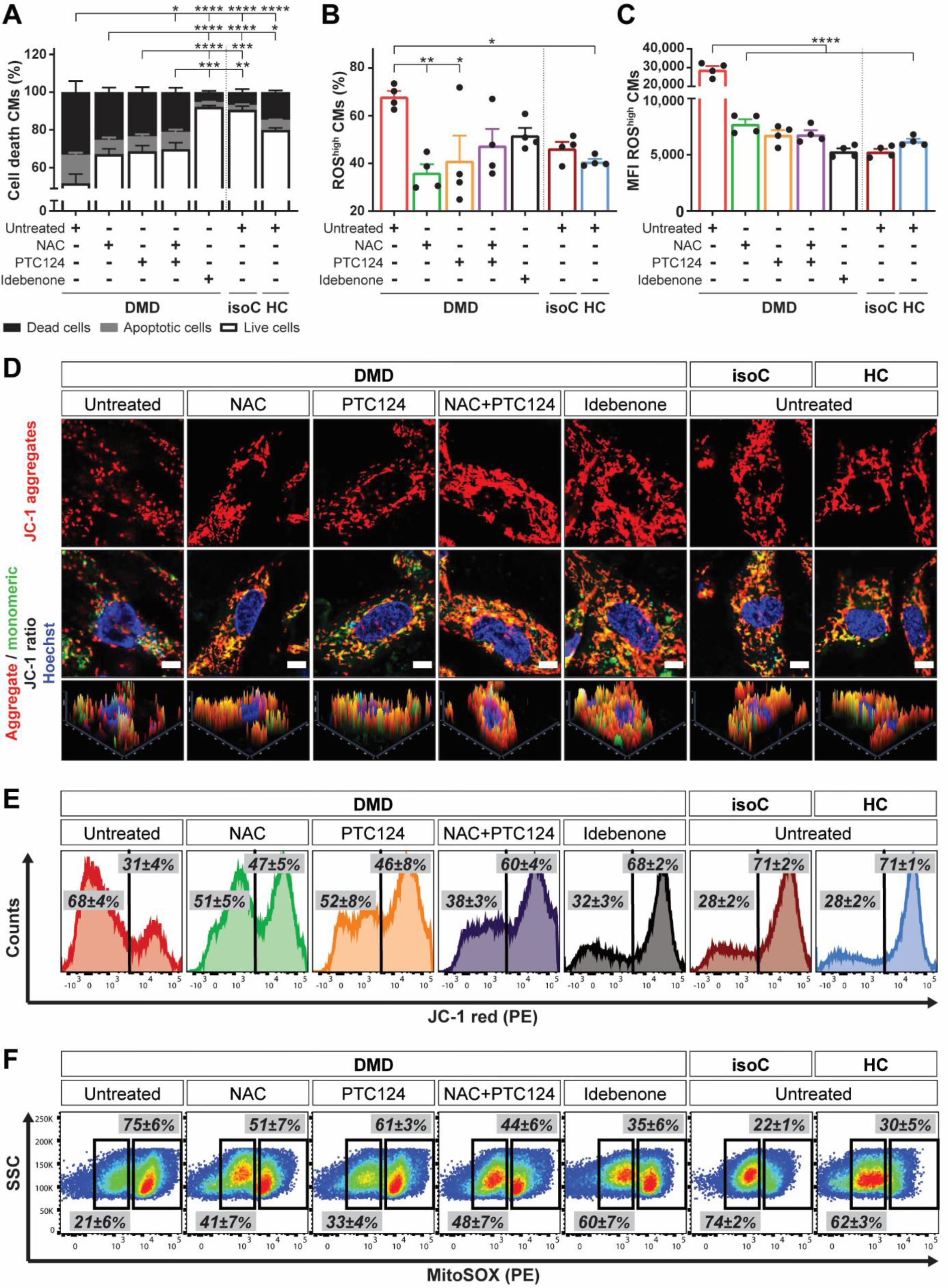
Characterization of the cardiomyopathic phenotype *in vitro* of DMD iPSC-CMs, showing premature cell death, depolarized mitochondria and increased intracellular ROS levels, which were counteracted by NAC, ataluren (PTC124) and idebenone. Flow cytometric quantification at day 15 of cardiac differentiation showing the percentage of cell death of Signal-Regulatory Protein Alpha (SIRPA) positive iPSC-CMs **(A)**, the percentage of CMs with high intracellular ROS levels **(B)** and the MFI of intracellular ROS in CMs **(C)** in conditions with (NAC, PTC124 and idebenone) or without (untreated) treatments. Data were representative of four independent experiments (N = 4) and values were reported as mean ± SEM. Significance of the difference was indicated as *P < 0.05; **P < 0.01; ***P < 0.001 and ****P < 0.0001. **(D)** Immunostaining of the fluorescent voltage-sensitive dye JC-1 was used to determine ΔΨm and mitochondrial health in 15-day-old differentiated iPSC-CMs. Untreated DMD iPSC-CMs were characterized by mitochondrial depolarization, as indicated by the decrease in mitochondrial aggregates (JC-1 red, *upper panels*) and the increase in mitochondrial monomers (JC-1 green, *middle panels*) in respect to treated DMD iPSC-CMs and controls. Corresponding histograms (*lower panels*) showed the JC-1 fluorescence intensity ratios (aggregates/monomers). Scale bar = 5 μm. **(E)** Representative flow cytometric analyses at day 15 of differentiation for JC-1 aggregates (PE) and JC-1 monomers (FITC) in DMD iPSC-CMs upon treatment. Data were representative of four independent experiments (N = 4). **(F)** Flow cytometric analyses at day 15 of differentiation showing the mitochondrial superoxide production (MitoSOX, PE) in depolarized DMD mitochondria compared to DMD isogenic and healthy controls. Data were representative of four independent experiments (N = 4). Flow cytometry data were reported as mean ± SEM.

### Dystrophin-Deficient iPSC-CMs Are Characterized by Depolarized Mitochondria

DMD pathology is accompanied by abnormal intracellular Ca^2+^ handling and the accumulation of dysfunctional mitochondria with defective structure (*31-38*). A distinctive feature of early phase cell death is the loss of the membrane potential of active mitochondria (ΔΨm) (*39*). Here, the carbocyanine compound JC-1, a fluorescent voltage-sensitive dye with membrane-permeant fluorescent lipophilic cationic properties (*40*), was used to determine ΔΨm in iPSC-CMs and mitochondrial health. Consistently with the previously observed accelerated death of untreated DMD iPSC-CMs, these cultures were characterized by mitochondrial depolarization, indicated by the decrease in red (aggregates)/green (monomers) JC-1 fluorescence intensity ratio (DMD, JC-1 aggregates: 31 ± 4% and JC-1 monomers: 68 ± 4%) compared to DMD isogenic (isoC, respectively 71 ± 2% and 28 ± 2%) and healthy controls (HC, respectively 71 ± 1% and 28 ± 2%; Fig. 3D-E and Fig. S6B). Interestingly, the combinatorial treatment of NAC and PTC124 (DMD NAC+PTC124, JC-1 aggregates: 60 ± 4% and JC-1 monomers: 38 ± 3%), as well as idebenone treatment (DMD idebenone, respectively 68 ± 2% and 32 ± 3%) displayed significantly beneficial effects on ΔΨm with respect to untreated DMD iPSC-CMs (DMD untreated, respectively 31 ± 4% and 68 ± 4%). Furthermore, flow cytometric analyses confirmed a significant increased superoxide production in depolarized mitochondria (DMD, 75 ± 6%) compared to controls (isoC, 22 ± 1% and HC, 30 ± 5%; Fig. 3F and Fig. S6C). No significant differences were observed for mitochondrial content upon the different treatments (Fig. S6D-E). The specificity of the drug effect on ΔΨm and on the mitochondrial superoxide concentrations of the experimental groups is shown in Supplemental Information (Fig. S8A-D). Taken together, these results indicate dysfunctional depolarized mitochondria in DMD iPSC-CMs, which could lead to excessive ROS leakage. The combined treatment of NAC and PTC124, as well of idebenone to DMD iPSC-CM cultures were able to rescue this condition.

### NOX4 Is Overexpressed in DMD iPSC-CMs

Several independent studies have reported increased NOX4 expression and activity in chronic heart failure, supporting the clinical relevance, although the role of NOX4 in CMs is still unclear (*12-16*). Here, the NOX2 and NOX4 isoforms, the predominantly expressed members of the ROS-producing NOX family in the heart, were investigated. Gene expression profiles did not reveal differential expression for *NOX2* and accessory regulatory subunits (*p47^phox^, p67^phox^,* and *RAC2* and *RAC3;* Fig. 4A). Interestingly, *NOX4* and its regulatory subunit *p22^phox^* were significantly upregulated in DMD iPSC-CMs. Moreover, DMD iPSC-CMs treated with PTC124 alone or in combination with NAC exhibited decreased *NOX4* and *p22^phox^* gene levels. In contrast, upon idebenone treatment, no reduction was observed in the expression of both genes. Flow cytometric analyses demonstrated a significant increased percentage of NOX4 positive DMD iPSC-CMs (DMD, 78 ± 3%) compared to DMD isogenic (isoC, 31 ± 4%) and healthy controls (HC, 29 ± 1%; Fig. 4B-C). The percentage of NOX4 positive DMD iPSC-CMs was reduced upon idebenone treatment (DMD idebenone, 34 ± 4% vs. isoC, 31 ± 4% and vs. HC, 29 ± 1%). The specificity of the drug on the expression of NOX4 among the experimental groups is shown in Supplemental Information (Fig. S9A-B). Western blot analysis confirmed significantly increased protein levels of NOX4 in DMD iPSC-CMs. (Fig. 4D). Upon idebenone addition, DMD iPSC-CMs showed downregulated NOX4 expression like observed in controls. These data demonstrate a significantly increased NOX4 expression in DMD iPSC-CMs that upon treatment with idebenone could be reverted to physiological levels.

**Fig. 4:**
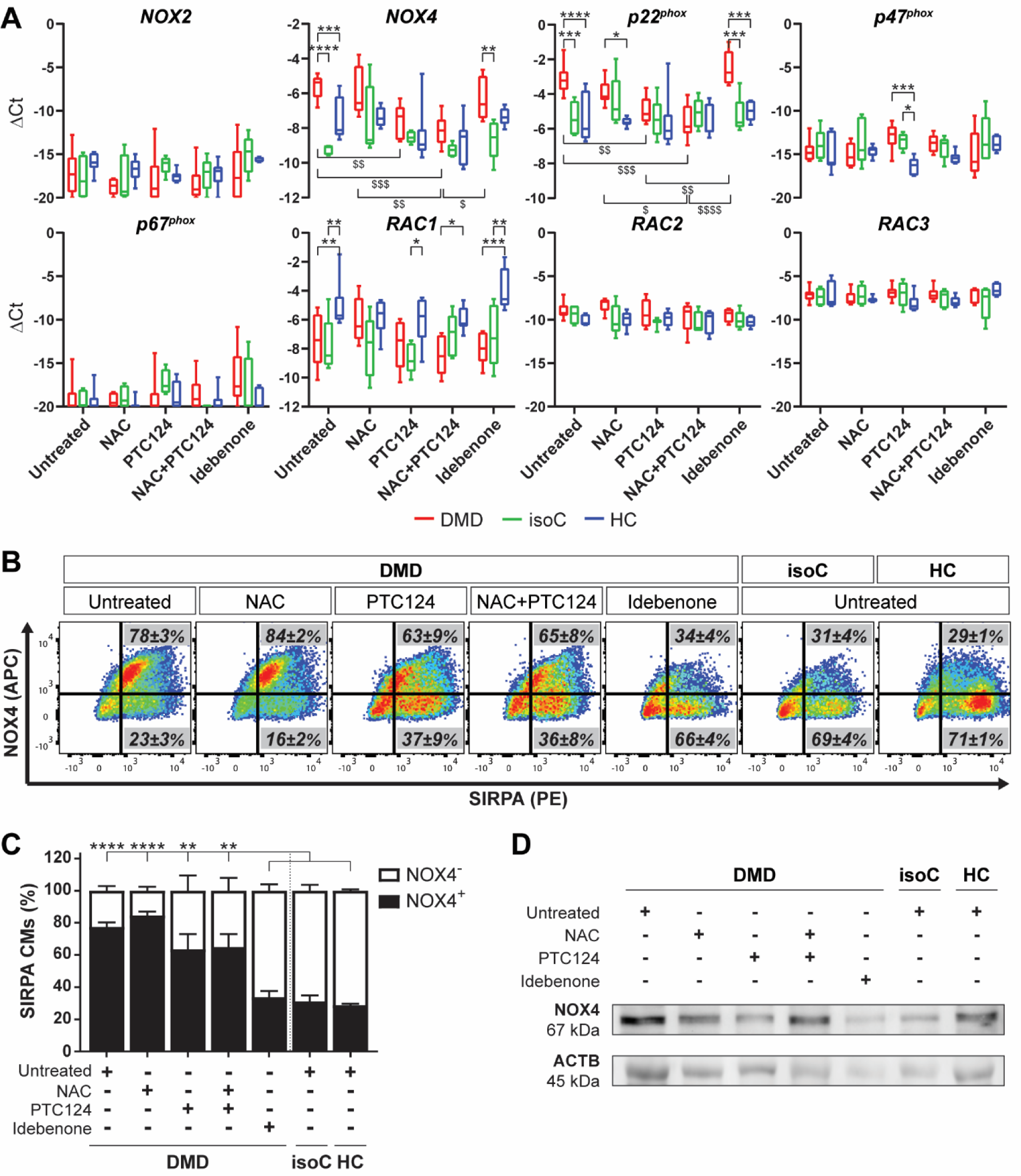
Increased expression levels of the ROS-producing NOX family enzyme NOX4 and its accessory regulatory subunit p22^phox^ in Dystrophin-Deficient iPSC-CM cultures. **(A)** Gene expression profiles at day 24 of cardiac differentiation of *NOX2* and *NOX4*, and the regulatory subunits (*p22^phox^, p47^phox^, p67^phox^, RAC1, RAC2* and *RAC3*) in DMD, DMD isogenic and healthy control iPSC-CMs upon treatment with NAC, PTC124 and idebenone. Each data point was represented as ΔCt, normalized for the housekeeping genes (*GAPDH* and *RPL13a*). Data were representative of five or more independent experiments (N ≥ 5) and values were expressed as mean ± SEM. Significance of the difference was indicated as follows: *P < 0.05; **P < 0.01; ***P < 0.001 and ****P < 0.0001 vs. subjects within the treatment condition; or ^$^P < 0.05; ^$$^P < 0.01; ^$$$^P < 0.001 and ^$$$$^P < 0.0001 vs. treatment conditions within the subject group. **(B)** Representative flow cytometric analyses at day 15 of differentiation showing the percentage of NOX4 (APC) protein expression in SIRPA (PE) positive DMD iPSC-CMs upon treatment. Data were representative of three independent experiments (N = 3). Flow cytometry data were reported as mean ± SEM. **(C)** Flow cytometric quantification at day 15 of differentiation of the percentage of SIRPA positive iPSC-CMs expressing NOX4 upon treatment. Data were representative of three independent experiments (N = 3) and values were expressed as mean ± SEM. Significance of the difference was indicated as follows: *P < 0.05; **P < 0.01; ***P < 0.001 and ****P < 0.0001. **(D)** Western blot analysis quantifying the protein expression levels of NOX4 in 15-day-old differentiated DMD and control iPSC-CMs, normalized to the loading protein ACTB.

**Fig. 5:**
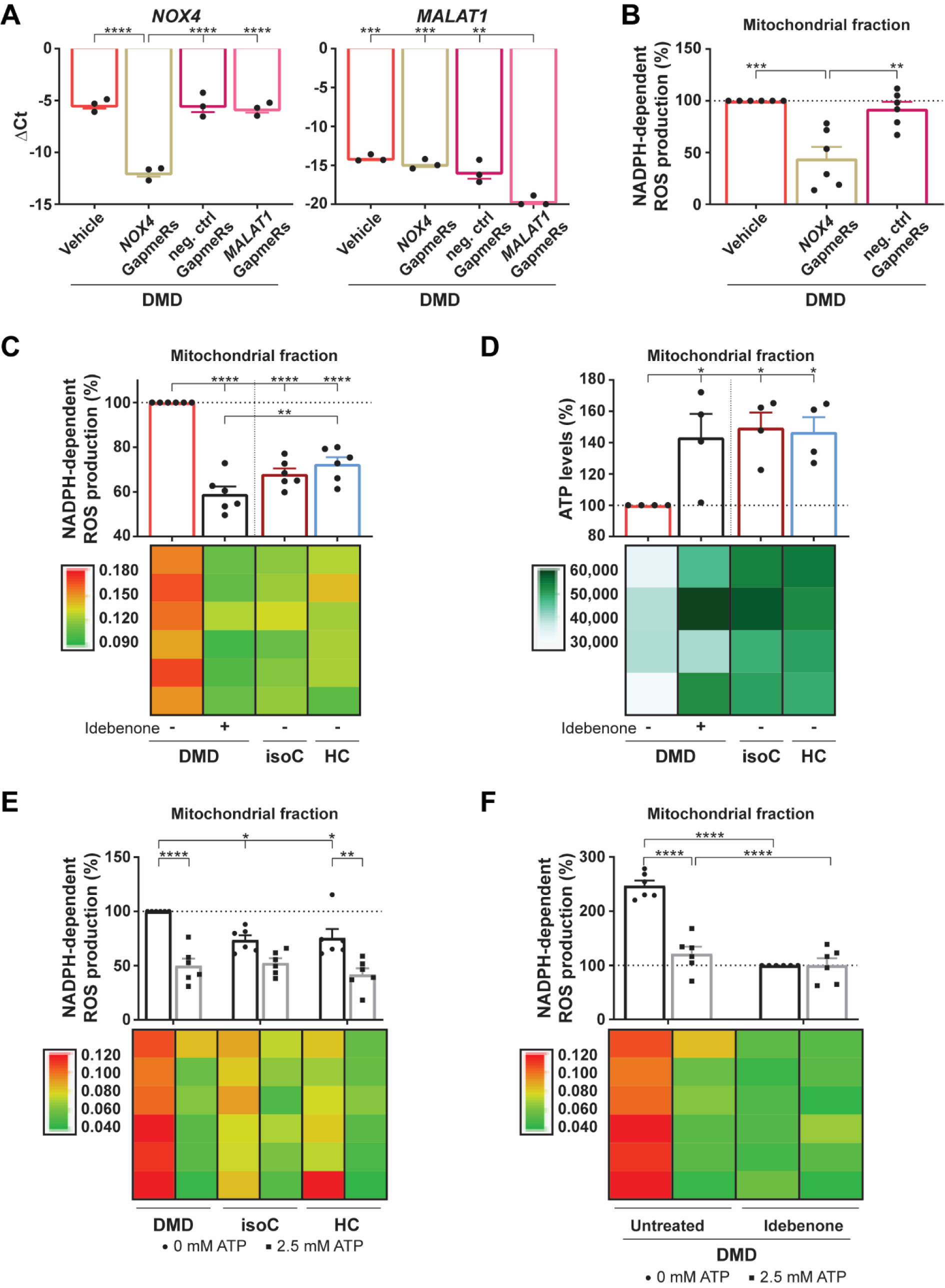
Idebenone could counteract the oxidative stress in DMD iPSC-CMs through ATP stimulation of the mitochondrial ETC, which, on its turn, reduced ROS-producing NOX4 activity. **(A)** Quantitative RT-PCR of *NOX4* gene expression levels after the addition of NOX4 targeted Antisense LNA GapmeRs to the DMD iPSC-CM cultures (*left panel*). As positive control for the efficiency of the Antisense LNA GapmeRs, *MALAT1* levels were determined after the addition of MALAT1 targeted Antisense LNA GapmeRs to the DMD iPSC-CM cultures (*right panel*). Each data point was represented as ΔCt, normalized for the housekeeping genes (*GAPDH* and *RPL13a*). Data were representative of three independent experiments (N = 3) and values were expressed as mean ± SEM. Significance of the difference was indicated as follows: *P < 0.05; **P < 0.01; ***P < 0.001 and ****P < 0.0001. **(B)** Quantification of the NOX4-mediated ROS production, measured via the NADPH-dependent ROS generation, in the isolated mitochondrial fraction of DMD iPSC-CMs after a 6 days preincubation with GapmeRs, inducing *NOX4* mRNA transient degradation. Each data point was represented as percentage (%), normalized to the mitochondrial fraction of the untreated DMD iPSC-CMs (vehicle). **(C)** Quantification of the NADPH-dependent ROS production of NOX4 in the mitochondrial fraction of DMD iPSC-CMs with or without idebenone treatment compared to DMD isogenic and healthy controls. **(D)** ATP luminescence detection showing the effect of idebenone treatment on the mitochondrial ATP levels in DMD iPSC-CMs. **(E)** Quantification of the ROS-producing NOX4 activity after 2.5 mM ATP addition in DMD iPSC-CM and control cultures. Each data point was represented as percentage (%), normalized to the mitochondrial fraction of the untreated DMD iPSC-CMs. **(F)** Quantification of the NADPH-dependent ROS production of NOX4 in the mitochondrial fraction of DMD iPSC-CMs upon 2.5 mM ATP addition, with or without idebenone treatment. Each data point was represented as percentage (%), normalized to the mitochondrial fraction of the idebenone-treated DMD iPSC-CM cultures. Data were representative of four or six independent experiments (N = 4 or N = 6) and values were expressed as mean ± SEM. Colored rectangles represented the independent experiments. Significance of the difference was indicated as follows: *P < 0.05; **P < 0.01; ***P < 0.001 and ****P < 0.0001.

Additionally, we demonstrated that the NOX4 upregulation in DMD iPSC-CMs (DMD #2 in Table S1) was not a common downstream pathway of cell death. Therefore, we preincubated iPSC-CMs with 1 μM STS for 6 h, a potent cell death inducer (*41*), and did not observe any increase in the NOX4 expression (Fig. S10A-B). Interestingly, by analyzing ΔΨm and the mitochondrial superoxide production in various DMD patient-specific iPSC-CM lines (DMD #2, DMD #5 and DMD #6 in Table S1), we could observe an association between the levels of mitochondrial depolarization and ROS production with the gene and protein levels of NOX4, suggesting a crucial role of NOX4 (Fig. S11A-F).

### Idebenone Stimulates ATP Production in Depolarized Mitochondria, Ameliorating NOX4-Mediated ROS Overproduction

Overall, oxidative stress, in synergy with intracellular Ca^2+^ overload, results in progressive worsening of DMD cardiomyopathy (*7, 37*). We hypothesized that *Dystrophin* gene mutations elicit excessive ROS generation via the mitochondrial ETC of dysfunctional mitochondria and a NOX4-based NADPH-dependent process. To assess whether increased NOX4 could contribute to elevated intracellular ROS concentrations, *NOX4* mRNA levels were transiently degraded by the addition of Antisense LNA GapmeRs to the DMD iPSC-CM cultures (Fig. 5A, *left panel*). Antisense LNA GapmeRs targeting *MALAT1* mRNA were used as positive control (Fig. 5A, *right panel*). Interestingly, transient GapmeR-induced *NOX4* mRNA degradation significantly reduced the NOX4 activity, monitored through changes in NADPH absorption (Fig. 5B) (*42, 43*). DMD iPSC-CMs exhibited significantly elevated NOX4 activity compared to controls (Fig. 5C). However, when idebenone was added to DMD iPSC-CMs, the NOX4 NADPH-dependent ROS production was significantly reduced in isolated mitochondria (Fig. 5C) and the total CM fraction (Fig. S12A). Moreover, idebenone restored ATP levels due to its electron donating property for mitochondrial ETC stimulation (Fig. 5D and Fig. S12B).

Recent studies have identified an ATP-binding motif within NOX4 through which ATP, upon binding, could regulate NOX4 activity (*43*). Adding dose-dependent ATP concentrations to DMD iPSC-CM cultures demonstrated that 2.5 mM ATP had a beneficial effect and significantly reduced ROS production of the NOX4 activity in respect to no ATP addition (Fig. 5E and Fig. S12C). Interestingly, idebenone alone or in combination with 2.5 mM ATP addition did ameliorate the activity of NOX4, in a similar manner, resulting in a significantly decreased NADPH-dependent ROS production compared to untreated DMD iPSC-CMs (Fig. 5F and Fig. S12D). The specificity of idebenone on the ROS-producing activity of NOX4 and on the ATP levels of the experimental groups is shown in Supplemental Information (Fig. S13A-D). These findings reveal an increased mitochondrial ROS-producing NOX4 activity in DMD iPSC-CMs, which was counteracted by idebenone application through ATP.

### DMD EHTs Show Improved Contractile Function after Idebenone Administration

In order to assess the amplitude of contraction of 3D EHT constructs, we monitored the micropost deflection movements of the EHT devices as a result of spontaneous contraction of the EHTs attached to the flexible microposts (Fig. 2B). At physiological 1.8 mM Ca^2+^ concentrations, the contractile function of untreated DMD iPSC-CM EHTs was significantly lower than untreated EHTs generated from isogenic or healthy iPSC-CMs (Fig. 6A), confirming the validity of the 3D EHT model system for DMD. However, DMD EHTs treated with idebenone exhibited a significantly increased contraction, whereas the combined treatment of idebenone and PTC124 improved even further the contractile function. By incubating DMD EHTs with various Ca^2+^ concentrations (ranging from 0.1 to 2.5 mM), we wanted to analyze the amplitude of contraction of DMD EHTs at physiological Ca^2+^ levels (1.8 mM; dotted line) and higher Ca^2+^ levels (2.5 mM), mimicking the detrimental increased Ca^2+^ environment, as reported in the heart from DMD patients (*35, 44, 45*). At physiological Ca^2+^ levels, idebenone had beneficial effects on the contraction of DMD EHTs compared to untreated DMD EHTs, whereas the contractile function of DMD EHTs did not show any improvements upon idebenone administration at 2.5 mM Ca^2+^ concentrations (Fig. 6B). However, the contractile function was significantly improved after the combinatorial treatment of idebenone and PTC124. These data point out the beneficial effect of a combinatorial treatment of idebenone and PTC124, highlighting the importance of targeting simultaneously different aspects of DMD cardiomyopathy in terms of heart functionality.

**Fig. 6:**
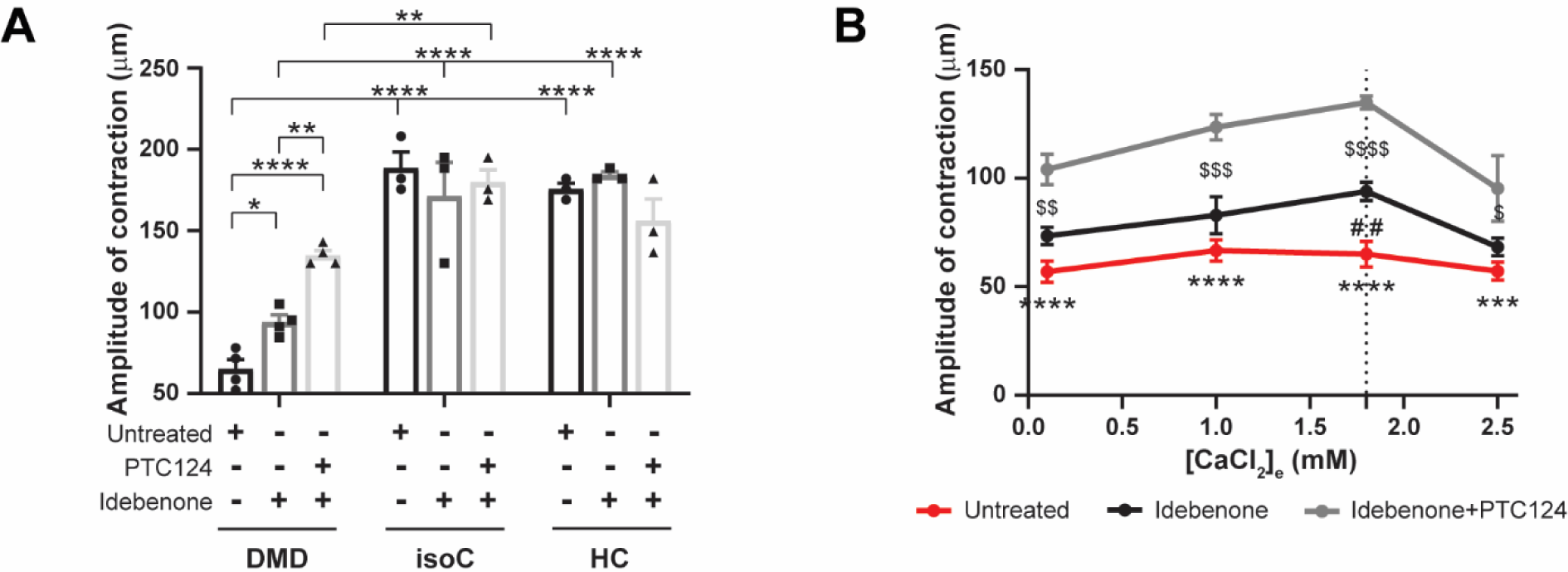
Improved contraction of 3D EHT constructs after administration of idebenone alone or idebenone in combination with PTC124 under physiological Ca^2+^ levels. **(A)** Spontaneous contraction and relaxation cycles of EHTs were monitored under temperature-controlled conditions (37°C) at 1.8 mM physiological Ca^2+^ concentrations, and measured by the deflection movements of the microposts (in μm). The effect of idebenone and PTC124 on the contractility of EHTs derived from DMD iPSC-CMs (EHT diameter in μm: 1,041.9 ± 74.1) was compared to DMD isogenic (diameter in μm: 938.0 ± 86.6) and healthy control EHTs (diameter in μm: 849.9 ± 80.5). Data were representative of three or four independent experiments (N ≥ 3) and values were expressed as mean ± SEM. Significance of the difference was indicated as follows: *P < 0.05; **P < 0.01; ***P < 0.001 and ****P < 0.0001. **(B)** EHTs derived from DMD iPSC-CMs were incubated with various Ca^2+^ concentrations (ranging from 0.1 to 2.5 mM) to assess the amplitude of contraction. Data were representative of three or four independent experiments (N ≥ 3) and values were expressed as mean ± SEM. Significance of the difference was indicated as follows: *P < 0.05; **P < 0.01; ***P < 0.001 and ****P < 0.0001 (untreated vs. idebenone+PTC124); or ^$^P < 0.05; ^$$^P < 0.01; ^$$$^P < 0.001 and ^$$$$^P < 0.0001 (idebenone vs. idebenone+PTC124); or ^#^P < 0.05; ^##^P < 0.01; ^###^P < 0.001 and ^####^P < 0.0001 (untreated vs. idebenone).

## DISCUSSION

Human iPSCs have the potential to differentiate in functional cell types that can be used as unlimited cell source of inaccessible tissues to study genetic disorders and, consequently, to gain novel insights in signaling pathways involved in the disease pathology.

In this study, we generated iPSC-based cardiac disease models from sample material of three DMD patients to study the early stages of cardiomyopathy in DMD. Human iPSCs were differentiated towards CMs according to the protocol of Burridge *et al.* (Nature Methods) (*29*) and Breckwoldt *et al.* (Nature Protocols) (*30*). First, human iPSCs were differentiated in a monolayer-based method using a fully chemically defined medium, consisting of the basal medium RPMI 1640, rice-derived recombinant human albumin and L-ascorbic acid 2-phosphate along with small molecule-based induction of differentiation (*29*). L-ascorbic acid 2-phosphate has been shown to enhance cardiac differentiation and maturation through increased collagen production by promoting cardiac progenitor cell proliferation via the MEK-ERK1/2 pathway. Furthermore, L-ascorbic acid 2-phosphate-induced CMs exhibited better sarcomeric organization and enhanced responses of APs and Ca^2+^ transients to β-adrenergic and muscarinic stimulations (*46*). Second, iPSC-CMs were further differentiated in 3D fibrin-based EHT constructs for contractility measurements (*30*). In several cancer-related studies the effect of ascorbic acid on ROS production has been reported (*47, 48*). In these studies, a ROS-scavenger effect was observed after the addition of 1 mM or higher concentrations of ascorbic acid. We used a lower final concentration, suggesting no significant antioxidative effect on ROS levels. Interestingly, Bartsch *et al.* (*49*) demonstrated an ascorbic acid enhanced cardiac differentiation accompanied by an upregulation of the NADPH oxidase isoforms NOX2 and NOX4 at basal expression levels with intracellular physiological ROS concentrations, indicating the suitability of the applied cardiac differentiation methods.

Human iPSC-CMs obtained from DMD patients represent hallmarks of DMD-associated heart complications in *in vitro* cultures. Published studies showed that the lack of Dystrophin in DMD iPSC-CMs resulted in enhanced cell death (*32*), Ca^2+^ handling abnormalities and reduced contractile function (*31, 33-36, 44, 45, 50*). We observed premature cell death of DMD iPSC-CMs due to significantly elevated intracellular oxidative stress levels. Furthermore, detailed characterization demonstrated mitochondrial depolarization and significantly increased NOX4 expression. Whether the abnormally upregulated NOX4 expression and its increased basal rate of ROS production are a direct or indirect consequence of the absence of Dystrophin is currently unknown. Increased Nox4 protein have been found in left ventricular CMs of *mdx* mice, associated with fibrosis and altered functional parameters in the heart (*14*). Deep RNA sequencing of the cardiac transcriptome on explanted human heart samples, obtained from patients suffering from heart failure, indicated extensive alternative splicing of the *NOX4* gene, associated with upregulation of the full-length NOX4 protein (*15*). In consistency with these results, we found a significantly increased expression and activity of the cardiac-specific ROS-producing NOX4 isoform in DMD iPSC-CMs. Dystrophin-deficient CMs are more vulnerable to mechanical stress due to the increased membrane fragility and stretch-induced Ca^2+^ influx, resulting in cell death (*32, 35, 44, 45*). The complexity of the DMD pathology results from the signal amplification systems, with bidirectional cross-talk and positive feed-back loops. ROS generation in response to mechanical forces may originate from diverse sources including mitochondria and NOX isoforms (*12, 13, 16*), or even other oxidase systems (*19, 51, 52*).

To ameliorate the DMD disease phenotype, we applied several therapeutic approaches. We investigated whether NAC, ataluren (PTC124) and idebenone could have beneficial effects on the dystrophic features observed in DMD iPSC-CM cultures. PTC124 drug efficacy analyses were performed only on the DMD iPSC line with the nonsense mutation in exon 35 (c.4,996C > T; p.Arg1,666X) of the *Dystrophin* gene. This line represents a subgroup of DMD patients (approximately 13%) that is responsive to the read-through chemical drug PTC124, which allowed us to investigate the effects of PTC124 on DMD cardiomyopathy in an *in vitro* iPSC-based disease model. PTC124 is one of the gene-based therapeutic approaches for DMD, although applicable for only a small subgroup of DMD patients with a nonsense mutation (*21*). We demonstrated re-expression of Dystrophin after PTC124 addition in a fraction of differentiating DMD iPSC-CMs. Recently, a phase 3 randomized placebo-controlled trial, evaluating an improvement in the 6-minute walking test after 48 weeks, has been completed (*53*), and a clinical trial to study Dystrophin expression levels in a small cohort of PTC124-treated patients with DMD is currently ongoing. These clinical studies aim at targeting the primary cause of DMD progression.

Nowadays several innovative therapeutic approaches focus on the secondary pathology. In the last decade, researchers have shown growing interest for idebenone as potential treatment for DMD. The precise mechanism by which idebenone exerts its protective effect is still unknown. Yet, idebenone has been reported to protect mitochondria from oxidative damage and boost their impaired function, delaying disease progression of DMD (*27, 28*). Interestingly, given the dual mode-of-action of idebenone (ROS-scavenger function and stimulation of the mitochondrial ETC), we showed that idebenone application exhibited a superior beneficial outcome on DMD iPSC-CMs through increased ATP production that, on its turn, decreased NOX4 activity. The exact mechanism of ATP-mediated inhibition of the NOX4 activity is still unclear.

Recently, an ATP-binding motif within the NOX4 isoform has been identified, suggesting a potential novel mechanism through which NOX4 can be allosterically regulated. During normal respiration, OXPHOS-driven ATP production in the mitochondria binds NOX4 through the ATP-binding domain, keeping the NOX4-produced ROS levels low (*43*). The ATP-binding motif (AXXXXGKT) (*54*) that resides within the amino acids 534-541 of the C-terminus, is unique to NOX4 (not found in other NOX isoforms) and is conserved in *Homo sapiens, Rattus norvegicus* and *Mus musculus* (*43*). In line with these results, we demonstrated that the addition of idebenone to DMD iPSC-CM cultures increased the intracellular and, more specifically, the mitochondrial ATP concentrations through idebenone-induced ETC stimulation. Moreover, idebenone could significantly reduce the ROS-producing NOX4 activity, assuming the allosterically regulation of NOX4 through ATP. Interestingly, the addition of external ATP to DMD iPSC-CM cultures resulted in a similar reduction of the NADPH-dependent ROS production of NOX4.

Elevated ATP concentrations can be used by skeletal and cardiac myosin to increase cross-bridge binding and cycling, leading to stronger and faster contraction and relaxation (*55*). Cardiac-specific overexpression by means of recombinant adeno-associated viral (rAAV)-mediated delivery of the enzyme ribonucleotide reductase that converts adenosine diphosphate (ADP) to deoxy-ADP (dADP), which, on its turn, is rapidly converted to deoxy-ATP (dATP) in cells, facilitated CM contraction and cardiac performance in normal rodent hearts as well as in rodent and pig infarcted hearts (*56, 57*). We showed improved contractile properties of EHTs derived from DMD iPSC-CMs upon idebenone administration at physiological Ca^2+^ concentrations. Preincubation of idebenone with PTC124 further enhanced the contractility, probably due to the PTC124-induced re-expression of Dystrophin proteins. In line with these results, the group of Olson performed CRISPR/Cas9-mediated exon skipping (“myoediting”) for DMD mutation corrections, in order to rescue the contractile dysfunction of DMD iPSC-CMs that were differentiated in 3D EHTs (*44, 45, 58*).

In conclusion, by using iPSC-CMs from DMD patients, we provided the first evidence that NOX4 expression and activity was significantly upregulated, contributing to high intracellular ROS and increased cell death. Furthermore, we compared the effects of the ROS scavenger NAC, the read-through premature termination codon chemical drug PTC124 and idebenone in an *in vitro* setting of cardiomyopathic DMD. Finally, we gained novel mechanistic insights in the mode-of-action of idebenone on the hyperactive state of NOX4-mediated ROS production. Idebenone-mediated stimulation of the ATP production by the ETC of mitochondria could increase the affinity of ATP to bind with NOX4, reducing the ROS-producing activity of NOX4. Considering the early cellular stress responses present in iPSC-CMs from DMD patients, interfering with any of these early cellular events that lead to excessive ROS signals would positively affect the mitochondrial activity resulting in an improved contractile function.

## MATERIALS AND METHODS

### Study Design

The objective of this study is to develop a stem cell-based model to investigate pathological mechanisms and evaluate their therapeutical potential in cardiomyopathy in DMD patients. The study was conducted in compliance with the principles of the Declaration of Helsinki, the principles of ‘Good Clinical Practice’ (GCP) and in accordance with all applicable regulatory requirements. The use of human samples from healthy control donors and DMD subjects for experimental purposes and protocols in the present study was approved by the Ethics Committee of the University Hospitals Leuven (respectively, S55438 and S65190). Subjects information, used in this study, is summarized in Table S1.

### Chemicals and Reagents

NAC (Merck), ataluren (PTC124; Selleckchem) and idebenone (Santhera Pharmaceuticals, Pratteln Switzerland). Staurosporine (STS; Merck). CM-H2DCFDA Total Intracellular ROS Indicator, JC-1 Mitochondrial Membrane Potential Probe, MitoSOX Red Mitochondrial Superoxide Indicator and MitoTracker-Red CMXRos Mitochondria Probe (all from Thermo Fisher Scientific). ATP Solution, Luminescent ATP Detection Assay Kit, Colorimetric NADPH Assay Kit (both from Abcam) and Mitochondrial Isolation Kit for Cultured Cells (Thermo Fisher Scientific).

### Generation of Integration-Free DMD iPSCs

hFs and hPBMCs were isolated from DMD patients with known *Dystrophin* mutations (Table S1). Somatic cells were reprogrammed towards pluripotency using the integration-free SeV-based technology, performed according to the manufacturer’s instructions (CytoTune-iPS 2.0 Sendai Reprogramming Kit; Thermo Fisher Scientific).

### Teratoma Formation Assay

Pluripotency of SeV-reprogrammed iPSCs was evaluated *in vivo* in 6- to 8-week-old immunodeficient *Rag2-null γc-null*/Balb/C mice. Teratoma formation experiments in mice were conducted following the guidelines of the Animal Welfare Committee of Leuven University and Belgian/European legislation (approved July 2016; P174/2016).

### Human iPSC Culture

Human control and DMD diseased iPSC lines (Table S1) were cultured feeder-free on Geltrex LDEV-Free hESC-Qualified Reduced Growth Factor Basement Membrane Matrix and maintained in Essential 8 Flex Basal Medium supplemented with Essential 8 Flex Supplement (50x) and 0.1% Pen/Strep (all from Thermo Fisher Scientific), at 37°C under normoxic conditions (21% O_2_ and 5% CO_2_). Colonies were routinely passaged non-enzymatically with 0.5 mM EDTA in Phosphate-Buffered Saline (PBS; both from Thermo Fisher Scientific). Mycoplasma contamination was assessed on a periodic basis for all cell cultures. No contaminated cells were used in the described experiments of this study.

### Generation of DMD Isogenic Control Line through CRISPR/Cas9 Genome Editing

To restore full-length expression of the *Dystrophin* gene, the isogenic control for the DMD iPSC patient line, characterized by a genetic point mutation in exon 35 (c.4,996C > T; p.Arg1,666X) of the *Dystrophin* gene, was generated through CRISPR/Cas9 from the *S. pyogenes* system (5’-NGG PAM) as previously described (*59*). Briefly, two 20-nucleotide sgRNAs (sgRNA #1: FW seq. CACCG-<u>ATTTAACCACTCTTCTGCTC</u> and RV seq. AAAC-<u>GAGCAGAAGAGTGGTTAAAT</u>-C; sgRNA #2: FW seq. CACCG-<u>TAACCACTCTTCTGCTCAGG</u> and RV seq. AAAC-<u>CCTGAGCAGAAGAGTGGTTA</u>-C) were designed and ligated into the RNA-guided nuclease plasmid (pX330-mCherry plasmid; Addgene), in order to induce the Cas9-mediated DSB in the genomic DNA of the Dystrophin-deficient iPSCs. Cas9-mediated genome editing was performed via HDR. The targeted DNA modification required the use of a plasmid-based donor repair template with two homology arm regions for the *Dystrophin* gene, flanking a GFP-Hygromycin-TK expressing cassette for selection. Here, one of the homology arms contained the genetic correction of the nonsense mutation in the *Dystrophin* gene. Finally, a completely gene editing-free DMD isogenic iPSC line was obtained due to PiggyBac excision and FIAU selection, restoring the expression of functional Dystrophin protein (Table S1).

### Monolayer-Based Cardiac Differentiation of Human iPSCs

Human iPSCs were differentiated into functional CMs according to a monolayer-based cardiac differentiation protocol, as previously described (*29*). Briefly, prior to differentiation, control and DMD iPSC lines were split in small colonies and subsequently cultured on a thin Matrigel Growth Factor Reduced (GFR) Basement Membrane Matrix layer (Corning) in complete Essential 8 Flex Medium at 37°C under hypoxic conditions (5% O_2_ and 5% CO_2_), in order to obtain the optimal confluency of 85%, three days after splitting. Mesoderm differentiation (day 0) was induced using 6 μM CHIR99021 (Axon Medchem) for 48 h in a chemically defined medium consisting of RPMI 1640 (Thermo Fisher Scientific), 500 μg/mL rice-derived recombinant human albumin and 213 μg/mL L-ascorbic acid 2-phosphate (both from Merck). After 24 h of CHIR99021 stimulation, iPSCs were transferred from hypoxia to normoxia. At day 2 of differentiation, iPSC-derived mesodermal cells were fed with basal medium supplemented with 4 μM IWR-1 (Merck) for 48 h, to induce cardiac progenitor cell differentiation. From day 4 onwards, medium was changed every other day with CM maintenance medium (RPMI 1640, rice-derived recombinant human albumin and L-ascorbic acid 2-phosphate). Contracting CMs appeared at day 8 or 9 of cardiac differentiation. DMD iPSC-CMs were treated with 3 mM NAC and 0.5 μM idebenone from day 8 onwards, and 20 μg/mL ataluren (PTC124) was supplemented to the cardiac differentiation medium from day 4 onwards. In *NOX4* knockdown experiments, 250 nM of single-stranded antisense oligonucleotides for silencing *NOX4* mRNA, called Antisense LNA GapmeRs (Qiagen), were added to the cell cultures at day 8 of differentiation.

### Generation of 3D EHT Constructs

3D EHT constructs were generated from 8- to 10-day-old iPSC-CMs, as previously described (*30*). CMs were dissociated with Collagenase A (1 U/mL; Merck) for 20 minutes at 37°C and transferred to custom-made 2% agarose (UltraPure; Thermo Fisher Scientific) casting molds in 24-well plate formats. The single-cell suspension was maintained in DMEM low glucose medium, containing 10% Fetal Bovine Serum (FBS), 1% heat-inactivated Horse Serum (HS), 1% Pen/Strep (all from Thermo Fisher Scientific) and 0.1% Rho-associated protein kinase (ROCK) inhibitor (Y-27632; VWR). Each EHT construct consisted of 1.0 x10^6^ cells, supplemented with GFR Matrigel, 5.06% fibrinogen (human plasma; Merck), 3U/mL thrombin (Stago BNL) and 1.44% aprotinin (Merck). The casting was performed around two flexible polydimethylsiloxane (PDMS) microposts within the agarose molds. After 2 h of incubation, polymerization formed a fibrin block around the microposts, embedding the single-cell suspension. The fibrin block was removed from the casting molds and transferred to 24-well plates, containing EHT medium composed of DMEM low glucose, 10% heat-inactivated HS, 1% Pen/Strep, 0.1% aprotinin and 0.1% Human Insulin Solution (Merck). Medium was changed every other day with EHT medium.

### Quantitative Real-Time PCR Analysis

Total RNA was extracted using the PureLink RNA Mini Kit and treated with the TURBO DNA-Free DNase Kit to assure highly pure RNA. 1 μg RNA was reverse transcribed into cDNA with SuperScript III Reverse Transcriptase First-Strand Synthesis SuperMix. Quantitative Real-Time PCR was performed with the Platinum SYBR Green qPCR SuperMix-UDG (all from Thermo Fisher Scientific). The oligonucleotide primer sequences (all from IDT) are listed in Table S2. A 10-fold dilution series ranging from 10^-3^ to 10^-8^ of 50 ng/μL human genomic DNA was used to evaluate the primer efficiency. Delta Ct (ΔCt) values were calculated by subtracting the Ct values from the genes of interest with the Ct values of the housekeeping genes (*GAPDH, HPRT* and *RPL13a*).

### Flow Cytometric Analysis

Differentiated iPSC-CMs were dissociated using Collagenase A (1 U/mL) for 20 minutes at 37°C. All flow cytometry procedures were performed according to the manufacturer’s instructions. Hank’s Balanced Salt Solution (HBSS; pH 7.2) with CaCl_2_ and MgCl_2_ supplemented with 2% FBS (both from Thermo Fisher Scientific), 10 mM HEPES and 10 mM NaN3 (both from Merck), was used as staining buffer. For high CM purity, iPSC-CMs were stained for the surface marker SIRPA (data not shown). If intracellular staining was necessary, cells were fixed with 4% paraformaldehyde (PFA; Polysciences) for 10 minutes at 37°C and permeabilized in ice-cold 90% methanol (Merck) for 30 minutes on ice, before the staining procedure. Fluorescence minus one (FMO) controls and compensations were included for appropriate gating. Samples were analyzed using the FACS Canto II HTS (BD Biosciences) and quantified using FlowJo Software Version 10 (FlowJo LLC). Table S3 provides a list of all flow cytometric antibodies used in this study.

### Immunofluorescence Imaging

Cells were fixed with 4% PFA for 10 minutes at 4°C, permeabilized for 30 minutes at room temperature in PBS supplemented with 0.2% Triton X-100 and 1% Bovine Serum Albumin (BSA) and blocked for 30 minutes at room temperature in 10% donkey serum (all from Merck). Samples were stained overnight at 4°C with the primary antibodies, followed by the appropriate secondary antibodies (1 h incubation at room temperature). Immunofluorescent primary and secondary antibodies were listed in Table S3. Nuclei were counterstained with 10 μg/mL Hoechst (33342; Thermo Fisher Scientific). Analyses were assessed using the Nikon Eclipse Ti Microscope or the Nikon Eclipse Ti A1R Configurated Confocal Microscope, with appropriate NIS-Elements Software (all from Nikon).

### Mitochondria and Cytoplasmic Fractionation

Mitochondrial and cytoplasmic separation was performed using the Mitochondrial Isolation Kit for Cultured Cells (Thermo Fisher Scientific), according to the manufacturer’s instructions with minor modifications. To obtain a more purified mitochondrial fraction (with a more than 50% reduction of the lysosomal and peroxisomal contaminants), the post-cell debris supernatant was subjected to an extra centrifuge step at 3000 x g for 15 minutes. For Western blot analysis, mitochondrial pellets were lysed with 2% CHAPS (Merck) in Tris-buffered saline (TBS; containing 25 mM Tris, 0.15 M NaCl; pH 7.2) and subsequently centrifuged at high speed for 2 minutes. Western blot analysis was performed on the supernatant, containing soluble mitochondrial protein.

### Western Blot Analysis

Western blot analysis for cell lysates was performed in RIPA buffer supplemented with 10 mM NaF, 0.5 mM Na3VO4, 1:100 protease inhibitor cocktail and 1 mM Phenylmethylsulfonyl Fluoride (PMSF; all from Merck). Equal amounts of protein (40 μg) were heat-denaturated at 95°C in sample-loading buffer (50 mM Tris-HCl, 100 mM DTT, 2% SDS, 0.1% bromophenol blue and 10% glycerol; pH 6.8), resolved by SDS-polyacrylamide gel electrophoresis and subsequently transferred to nitrocellulose membranes (Amersham Protran Western Blotting Membranes; Merck). The filters were blocked with TBS containing 0.05% Tween and 5% non-fat dry milk (Merck). Incubation was done overnight with the indicated primary antibody dilutions, as listed in Table S2. Horseradish peroxidase-conjugated secondary antibodies (Bio-Rad) were diluted 1:5,000 in TBS-Tween (0.05%) with 2.5% non-fat dry milk. After incubation with SuperSignal Pico or Femto chemiluminescence substrate (both from Thermo Fisher Scientific), the polypeptide bands were detected with GelDoc Chemiluminescence Detection System (Bio-Rad). Quantification of relative densitometry was obtained by normalizing to the background and to loading control proteins (ACTB, from Cell Signaling Technology) using Image Lab Software (Bio-Rad).

### Patch-Clamp Electrophysiology and Ca^2+^ Recordings

Single cells were seeded on Matrigel-coated coverslips. Cells were perfused at 37°C with a solution containing the following (in mM): 137 NaCl, 5.4 KCl, 1.8 CaCl_2_, 0.5 MgCl_2_, 10 glucose and 10 Na-HEPES. The pH was adjusted to 7.4 with NaOH. The patch-clamp pipettes were filled with a solution containing the following (in mM): 120 K-Asp, 20 KCl, 10 HEPES, 5 Mg-ATP, 10 NaCl and 0.05 K5Fluo-4. The pH was adjusted to 7.2 with KOH. Patch electrode resistances were between 2.5 and 3 MΩ when the pipettes were filled with intracellular solution. Cells were patched in the whole-cell configuration. Data were recorded using an Axopatch 200B amplifier (Axon Instruments) at a sampling rate of 10 kHz. Signals were filtered with 5 kHz low-pass Bessel filters. Action potentials (APs) were recorded in current-clamp mode, and if not spontaneous, after a 5 ms pulse of 0.5 nA at a 1 Hz frequency. Ca^2+^ currents were measured in voltage-clamp mode. After a Na^+^ current inactivation step from -70 mV to 40 mV for 750 ms, Ca^2+^ currents were recorded with 10 mV voltage steps from -40 mV to 60 mV during 205 ms. For analysis, the maximum amplitude of the Ca^2+^ current was measured and corrected for the cell capacitance. Data were analyzed with Clampfit Software (Axon Instruments).

### Contractility Measurements of 3D EHT Constructs

The contractile properties of 3D EHTs were monitored by measuring the deflection distances of the microposts of the EHT device (in μm) during spontaneous contraction and relaxation under temperature-controlled conditions (37°C) in oxygenated Tyrode’s solution (in mM; containing 137 NaCl, 5.4 KCl, 0.5 MgCl_2_, 12.8 HEPES and 5.5 Glucose; dissolved in deionized sterile water at pH 7.4) with Ca^2+^. A Ca^2+^ concentration of 1.8 mM was used to mimic physiological conditions. EHT constructs for contractility measurements were generated from 8-day-old iPSC-CMs and monitored after 5 days of EHT maturation.

### ATP Luminescence and NADPH Detection

The level of ATP was measured using the Luminescent ATP Detection Assay Kit (Abcam), according to the manufacturer’s instructions. Recordings were performed with the EG&G Berthold Microplate Luminometer LB 96V and corresponding software (Berthold Technologies). Using the NADPH Assay Kit (Abcam), NADPH-dependent ROS production was measured in the presence or absence of 2.5 or 5.0 mM ATP (preincubated for 60 minutes) in the total CM fraction or isolated mitochondrial fraction, according to the manufacturer’s instructions. Recordings were performed with the ELx808 Absorbance Microplate Reader and quantified using Gen5 Software Version 3 (both from BioTek Instruments).

### Statistical Analysis

Data were statistically analyzed using Prism Software Version 8 (GraphPad). All data were reported as mean ± standard error of the mean (SEM). Differences between two groups were examined for statistical significance using Student’s t-test. One-Way or Two-Way ANOVA (with multiple comparisons test and Tukey’s or Bonferroni’s correction) were used for three or more groups. Significance of the difference was indicated as follows: *P < 0.05; **P < 0.01; ***P < 0.001 and ****P < 0.0001.

## ACKNOWLEDGEMENTS

The authors thank Santhera Pharmaceuticals (Pratteln, Switzerland) for providing idebenone. The authors gratefully acknowledge Hanne Grosemans and Sylvia Sauvage for technical assistance; Ewa Berlińska, Filippo Conti, Vittoria Marini and Ilaria Tortorella for the assistance during their internship period at the KU Leuven; and Christina Vochten and Vicky Raets for the administrative assistance.

## Author contributions

R.D. participated in conception and design, collection and assembly of data, data analysis and interpretation, and manuscript writing. D.C. participated in western blot collection and assembly of data, data analysis and interpretation, and reviewed the manuscript. G.G. participated in patch-clamp electrophysiology and Ca^2+^ recordings, data analysis and interpretation, and reviewed the manuscript. L.D.W. provided DMD patient study samples, participated in conception and design, data analysis and interpretation, and reviewed the manuscript. N.G. provided DMD patient study samples, and reviewed the manuscript. K.D. participated in data analysis and interpretation, and reviewed the manuscript. C.M.V. participated in conception and design, data analysis and interpretation, and reviewed the manuscript. K.R.S. participated in conception and design, data analysis and interpretation, and reviewed the manuscript. G.M.B. and M.S. participated in conception and design, data analysis and interpretation, reviewed the manuscript, and gave final approval for manuscript submission.

## Data and materials availability

All data generated or analyzed during this study are included in this published article (and its supplementary information files).

## Supplementary Materials

**Table S1.** Characteristics of DMD subjects and iPSC lines.

**Table S2.** List of primers for Quantitative Real-Time PCR.

**Table S3.** List of antibodies for flow cytometry (FC), immunostaining (IF) and western blot (WB).

**Fig. S1.** Characterization of the DMD patient hF-iPSC clones, harboring the nonsense mutation in exon 35 (c.4,996C > T; p.Arg1,666X) of the *Dystrophin* gene, generated by the non-integrating SeV-mediated reprogramming method.

**Fig. S2.** Characterization of the pluripotency state of the CRISPR/Cas9 corrected DMD isogenic control line.

**Fig. S3.** Validation of the Cas9 cutting efficiency and analysis of the off-targets.

**Fig. S4.** Lack of Dystrophin protein in iPSC-CMs from DMD patients resulted in premature cell death.

**Fig. S5.** Dystrophin re-expression in DMD iPSC-CMs after PTC124 treatment.

**Fig. S6.** Corresponding flow cytometric graphs and quantification for the characterization of the cardiomyopathic phenotype of DMD iPSC-CMs, showing premature cell death, depolarized mitochondria and increased intracellular ROS levels.

**Fig. S7.** Drug specificity on cell death and intracellular ROS concentrations in the experimental iPSC-CM groups.

**Fig. S8.** Drug specificity on ΔΨm and mitochondrial superoxide concentrations in the experimental iPSC-CM groups.

**Fig. S9.** The specificity of the treatment options on the expression levels of NOX4 in iPSC-CM cultures.

**Fig. S10.** NOX4 protein expression levels after STS-induced cell death in DMD and control iPSC-CMs.

**Fig. S11.** Characterization of DMD patient-specific iPSC-CMs *in vitro*, showing increased NOX4 gene and protein expression levels, depolarized mitochondria and increased intracellular ROS levels.

**Fig. S12.** NADPH-dependent ROS production and intracellular ATP levels in DMD iPSC-CMs after idebenone application.

**Fig. S13.** The specificity of idebenone on the NADPH-dependent ROS production and ATP levels in the experimental iPSC-CM groups.

