## Supplementary Materials for "Human iPSC-Based Model Reveals NOX4 as Therapeutic Target in Duchenne Cardiomyopathy"

(Short Title: Overexpression of NOX4 in DMD CMs induced cell death)

Robin Duelen[1], Domiziana Costamagna[1,2], Guillaume Gilbert[3], Liesbeth De Waele[4], Nathalie Goemans[4], Kaat Desloovere[2], Catherine M. Verfaillie[5], Karin R. Sipido[3], Gunnar M. Buyse[4] and Maurilio Sampaolesi[1,6,*]

[1]*Translational Cardiomyology, Stem Cell and Developmental Biology, Dept. of Development and Regeneration, KU Leuven, 3000 Leuven, Belgium*

[2]*Research Group for Neurorehabilitation, Dept. of Rehabilitation Sciences, KU Leuven, 3000 Leuven, Belgium*

[3]*Experimental Cardiology, Dept. of Cardiovascular Sciences, KU Leuven, 3000 Leuven, Belgium*

[4]*Pediatric Neurology, University Hospitals Leuven, Dept. of Development and Regeneration, KU Leuven, 3000 Leuven, Belgium*

[5]*Stem Cell Institute Leuven, Stem Cell and Developmental Biology, Dept. of Development and Regeneration, KU Leuven, 3000 Leuven, Belgium*

[6]*Division of Human Anatomy, Dept. of Public Health, Experimental and Forensic Medicine, University of Pavia, 27100 Pavia, Italy*

[*]Corresponding author: Prof. Dr. Maurilio Sampaolesi

Translational Cardiomyology Lab

KU Leuven, Herestraat 49 – O&N4 – bus 804

3000 Leuven, Belgium

### SUPPLEMENTARY MATERIALS

**Table S1:** Characteristics of DMD subjects and iPSC lines.

| | ID | DMD #2 | DMD #5 | DMD #6 | DMD #2 isogenic | HC #1 | HC #2 | HC #3 |
|---|---|---|---|---|---|---|---|---|
| | **Origin** | hFs | hPBMCs | hPBMCs | DMD #2 | hPBMCs | hFs | hFs |
| | **Mutation** | pt.mut. exon 35 (c.4,996C > T; p.Arg1,666X) | del. exon 51-55 | del. exon 46-51 | CRISPR/Cas9 correction pt.mut. exon 35 | NA | NA | NA |
| | **Category** | diseased | diseased | diseased | healthy | healthy | healthy | healthy |
| **Phenotype** | **Age start steroids** | 7.5 years | 4.5 years | 6.5 years | NA | NA | NA | NA |
| | **Age loss of ambulation** | 8.5 years | 10 years | 11 years | | | | |
| **Cardiomyopathy** | **FS < 30%** | yes, 29% | no, 32% | no, 32% | | | | |
| | **EF** | 50% | 66% | 61% | | | | |
| | **Measured at age** | 20 years | 9 years | 20 years | | | | |
| | **Age at onset** | 10 years | NA | NA | | | | |
| **Pulmonary function** | **FVC < 50%** | yes, 7% | no, 80% | yes, 19% | | | | |
| | **Measured at age** | 25 years | 9 years | 19 years | | | | |

All DMD and control lines were previously characterized in-house or already published.

In the current study, somatic cells from DMD subjects were used to generate three human diseased iPSC lines DMD #2, DMD #5 and DMD #6 *(60)*. Four different human control lines were used: (1) the DMD isogenic control line was in-house generated through CRISPR/Cas9 gene editing, as described in Materials and Methods; (2) HC #1 is commercially available from Thermo Fisher Scientific (Catalog number A18945); (3) HC #2 was kindly provided by Prof. C. Verfaillie (University of Leuven, Belgium) and generated by transduction of the new-born male fibroblast BJ1 cell line, as published by Coll *et al. (61)*; and (4) HC #3 was a gift from Prof. P. Jennings (Medizinische Universität Innsbruck, Austria) to Prof. C. Verfaillie and generated by SeV-based reprogramming of male donor fibroblasts (SBAD2), as published by Rauch *et al. (62)*.

DMD: Duchenne muscular dystrophy; HC: healthy control; hPBMCs: human peripheral blood mononuclear cells; hFs: human fibroblasts; FS: fractional shortening; EF: ejection fraction; FVC: forced vital capacity; NA: not applicable.

**Table S2:** List of primers for Quantitative Real-Time PCR.

| Gene | Primer direction | Primer sequence (5’ > 3’) |
|---|---|---|
| ***c-MYC*** | forward | TCCTCGGATTCTCTGCTCTCCT |
| | reverse | AGAAGGTGATCCAGACTCTGACCT |
| ***Dystrophin*** | forward | ATGCTTTGGTGGGAAGAAGT |
| | reverse | GGGCATGAACTCTTGTGGAT |
| ***GAPDH*** | forward | TCAAGAAGGTGGTGAAGCAGG |
| | reverse | ACCAGGAAATGAGCTTGACAAA |
| ***GDF-3*** | forward | ACACCTGTGCCAGACTAAGATGCT |
| | reverse | TGACGGTGGCAGAGGTTCTTACAA |
| ***HPRT*** | forward | TGACACTGGCAAAACAATGCA |
| | reverse | GGTCCTTTTCACCAGCAAGCT |
| ***hTERT*** | forward | AAATGCGGCCCCTGTTTCT |
| | reverse | CAGTGCGTCTTGAGGAGCA |
| ***KLF-4*** | forward | CGGACATCAACGACGTGAG |
| | reverse | GACGCCTTCAGCACGAACT |
| ***MALAT1*** | forward | GGACTTGCCTCAACTCCCTC |
| | reverse | GCCCTCTCAGCCACTCAAAT |
| ***MYH6*** | forward | GCCCTTTGACATTCGCACTG |
| | reverse | CGGGACAAAATCTTGGCTTTGA |
| ***MYH7*** | forward | ACTGCCGAGACCGAGTATG |
| | reverse | GCGATCCTTGAGGTTGTAGAGC |
| ***MYL2*** | forward | TTGGGCGAGTGAACGTGAAAA |
| | reverse | CCGAACGTAATCAGCCTTCAG |
| ***MYL7*** | forward | ACATCATCACCCACGGAGAAGAGA |
| | reverse | ATTGGAACATGGCCTCTGGATGGA |
| ***NANOG*** | forward | TGGCCGAAGAATAGCAATGGTGTG |
| | reverse | TTCCAGGTCTGGTTGCTCCACATT |
| ***NOX2*** | forward | TGCCAGTCTGTCGAAATCTGC |
| | reverse | ACTCGGGCATTCACACACC |
| ***NOX4*** | forward | TCCGGAGCAATAAGCCAGTC |
| | reverse | CCATTCGGATTTCCATGACAT |
| ***OCT4*** | forward | CGAGCAATTTGCCAAGCTCCTGAA |
| | reverse | GCCGCAGCTTACACATGTTCTTGA |

| | | |
|---|---|---|
| ***p22***$^{phox}$ | forward | TACTATGTTCGGGCCGTCCT |
| | reverse | CACAGCCGCCAGTAGGTA |
| ***p47***$^{phox}$ | forward | GGGGCGATCAATCCAGAGAAC |
| | reverse | GTACTCGGTAAGTGTGCCCTG |
| ***p67***$^{phox}$ | forward | CCAGAAGCATTAACCGAGACAA |
| | reverse | CCTCGAAGCTGAATCAAGGC |
| ***RAC1*** | forward | ATGTCCGTGCAAAGTGGTATC |
| | reverse | CTCGGATCGCTTCGTCAAACA |
| ***RAC2*** | forward | TCTGCTTCTCCCTCGTCAG |
| | reverse | TCACCGAGTCAATCTCCTTGG |
| ***RAC3*** | forward | CTTCGAGAATGTTCGTGCCAA |
| | reverse | CCGCTCAATGGTGTCCTTG |
| ***REX1*** | forward | TGGAGGAATACCTGGCATTGACCT |
| | reverse | AGCGATTGCGCTCAGACTGTCATA |
| ***RPL13a*** | forward | CCTGGAGGAGAAGAGGAAAGAGA |
| | reverse | TTGAGGACCTCTGTGTATTTGTCAA |
| ***SOX2*** | forward | TGGCGAACCATCTCTGTGGT |
| | reverse | CCAACGGTGTCAACCTGCAT |
| ***TNNI1*** | forward | CCCAGCTCCACGAGGACTGAACA |
| | reverse | TTTGCGGGAGGCAGTGATCTTGG |
| ***TNNI3*** | forward | GATGCGGCTAGGGAACCTC |
| | reverse | GCATAAGCGCGGTAGTTGGA |

**Table S3:** List of antibodies for flow cytometry (FC), immunostaining (IF) and western blot (WB).

| **Protein** | **Antibody name** | **Provider** | **FC** | **IF** | **WB** |
|---|---|---|---|---|---|
| **ACTA2** | Anti-Alpha Smooth Muscle Actin (Mouse monoclonal) | Merck | | 1:300 | |
| **ACTB** | Beta Actin (13E5) (Rabbit monoclonal) | Cell Signaling Technology | | | 1:1,000 |
| **ACTN2** | Anti-Sarcomeric Alpha Actinin (EA-53) (Mouse monoclonal) | Abcam | | | 1:100 |
| **AFP** | Anti-Human Alpha-1-Fetoprotein (Rabbit polyclonal) | Dako | | 1:150 | |
| **Annexin V** | APC Annexin V | BD Pharmingen | 1:20 | | |
| **CASP3 (cleaved)** | Cleaved Caspase-3 (Asp175) (Rabbit polyclonal) | Cell Signaling Technology | | | 1:500 |
| **CASP3 (full length)** | Caspase-3 (Rabbit polyclonal) | Cell Signaling Technology | | | 1:1,000 |
| **cTnT** | Recombinant Anti-Cardiac Troponin T (EPR3696) (Rabbit monoclonal) | Abcam | | 1:100 | |
| **$p22^{phox}$** | Anti-Cytochrome b245 Light Chain/ p22-phox (44.1) (Mouse monoclonal) | Abcam | | | 1:100 |
| **Dystrophin (DMD)** | DYS1 (Rod Domain) (Mouse monoclonal) | Leica Novocastra | | 1:25 | |
| **Dystrophin (DMD)** | DYS2 (C-terminus) (Mouse monoclonal) | Leica Novocastra | | 1:25 | |
| **Dystrophin (DMD)** | DYS3 (N-terminus) (Mouse monoclonal) | Leica Novocastra | | 1:25 | |
| **LIN28** | LIN-28 (S-15) (Goat polyclonal) | Santa Cruz Biotechnology | | 1:50 | |

| | | | | | |
|---|---|---|---|---|---|
| **NANOG** | Nanog<br>(Rabbit polyclonal) | Thermo Fisher Scientific | | 1:200 | |
| **NOX4** | Anti-NADPH Oxidase 4<br>(Rabbit monoclonal) | Abcam | 1:2,175 | | 1:1,000 |
| **OCT4** | Anti-Oct4 - Embryonic Stem Cell Marker<br>(Goat polyclonal) | Abcam | | 1:200 | |
| **PARP** | PARP (46D11)<br>(Rabbit monoclonal) | Cell Signaling Technology | | | 1:1,000 |
| **SIRPA** | PE Anti-Human CD172a/b (SIRPα/β) | BioLegend | 1:100 | | |
| **SOX2** | Sox-2 (Y-17)<br>(Goat polyclonal) | Santa Cruz Biotechnology | | 1:50 | |
| **SSEA4** | SSEA-4 (MC813)<br>(Mouse monoclonal) | Santa Cruz Biotechnology | | 1:50 | |
| **TRA-1-60** | TRA-1-60 (TRA-1-60)<br>(Mouse monoclonal) | Santa Cruz Biotechnology | | 1:50 | |
| **TUBB** | Beta Tubulin<br>(Rabbit monoclonal) | NovusBio | | 1:250 | |

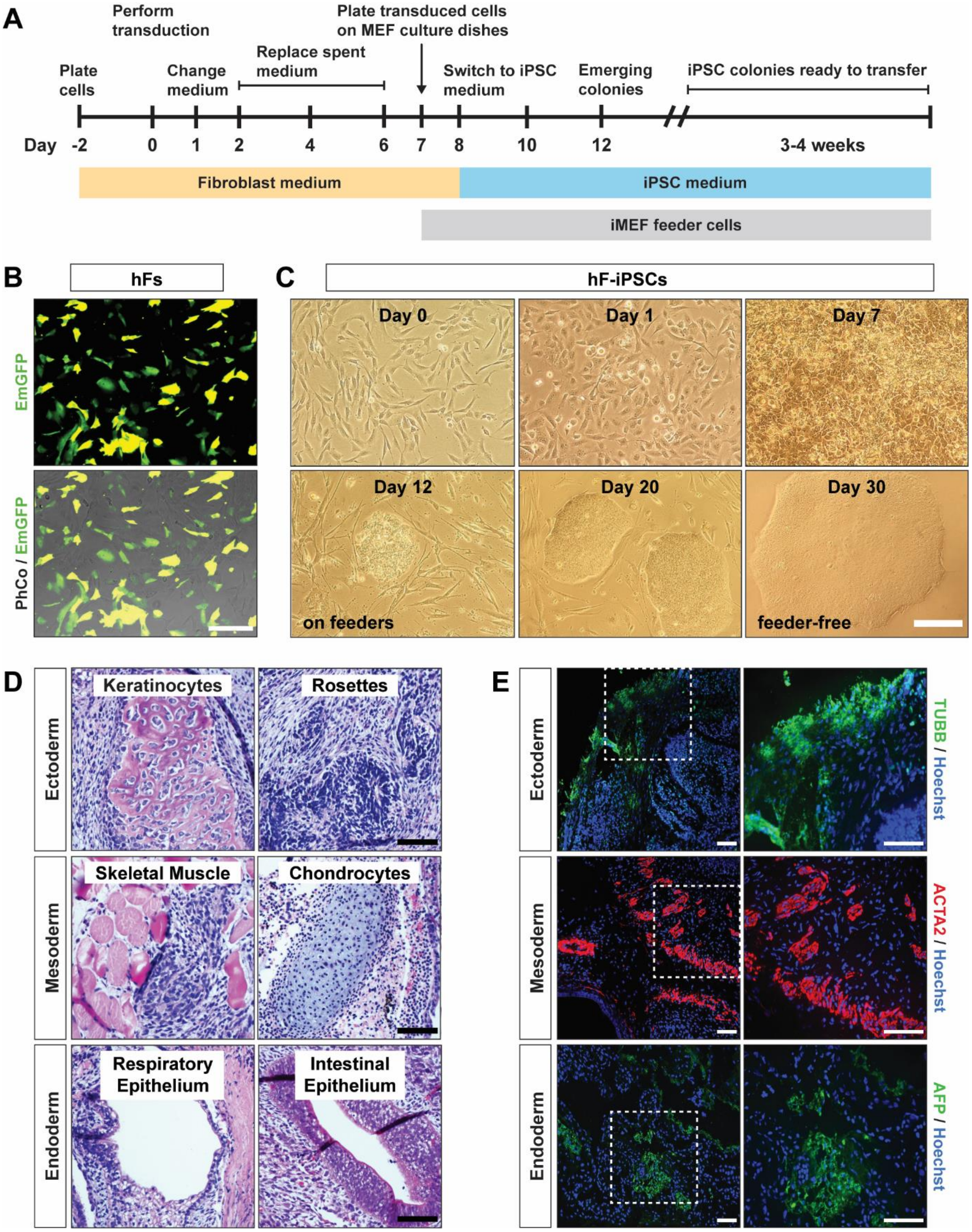


**Fig. S1: Characterization of the DMD patient hF-iPSC clones, harboring the nonsense mutation in exon 35 (c.4,996C > T; p.Arg1,666X) of the *Dystrophin* gene, generated by the non-integrating SeV-mediated reprogramming method. (A)** Schematic representation of the SeV iPSC reprogramming protocol for hFs (data not shown for hPBMCs). hFs were transduced at day 0 using the integration-free SeV vectors, expressing the OSKM (OCT3/4, SOX2, KLF4 and c-MYC) pluripotency markers. **(B)** EmGFP (green) expression of transduced DMD somatic cells, 1 day after transduction with SeV reprogramming vectors. **(C)** Morphological progression of DMD hFs towards iPSC clones. **(D)** Hematoxylin and eosin staining on SeV-reprogrammed iPSC-induced *in vivo* teratomas showing the derivatives of the three developmental germ layers, including keratinocytes and rosettes (ectoderm), skeletal muscle fibers and chondrocytes (mesoderm), and epithelium from respiratory and intestinal tract (endoderm). **(E)** Immunostaining of three germ lineage markers: Beta Tubulin (TUBB; ectoderm), Alpha Smooth Muscle Actin (ACTA2; mesoderm) and Alpha Fetoprotein (AFP; endoderm). Scale bar = 100 μm.

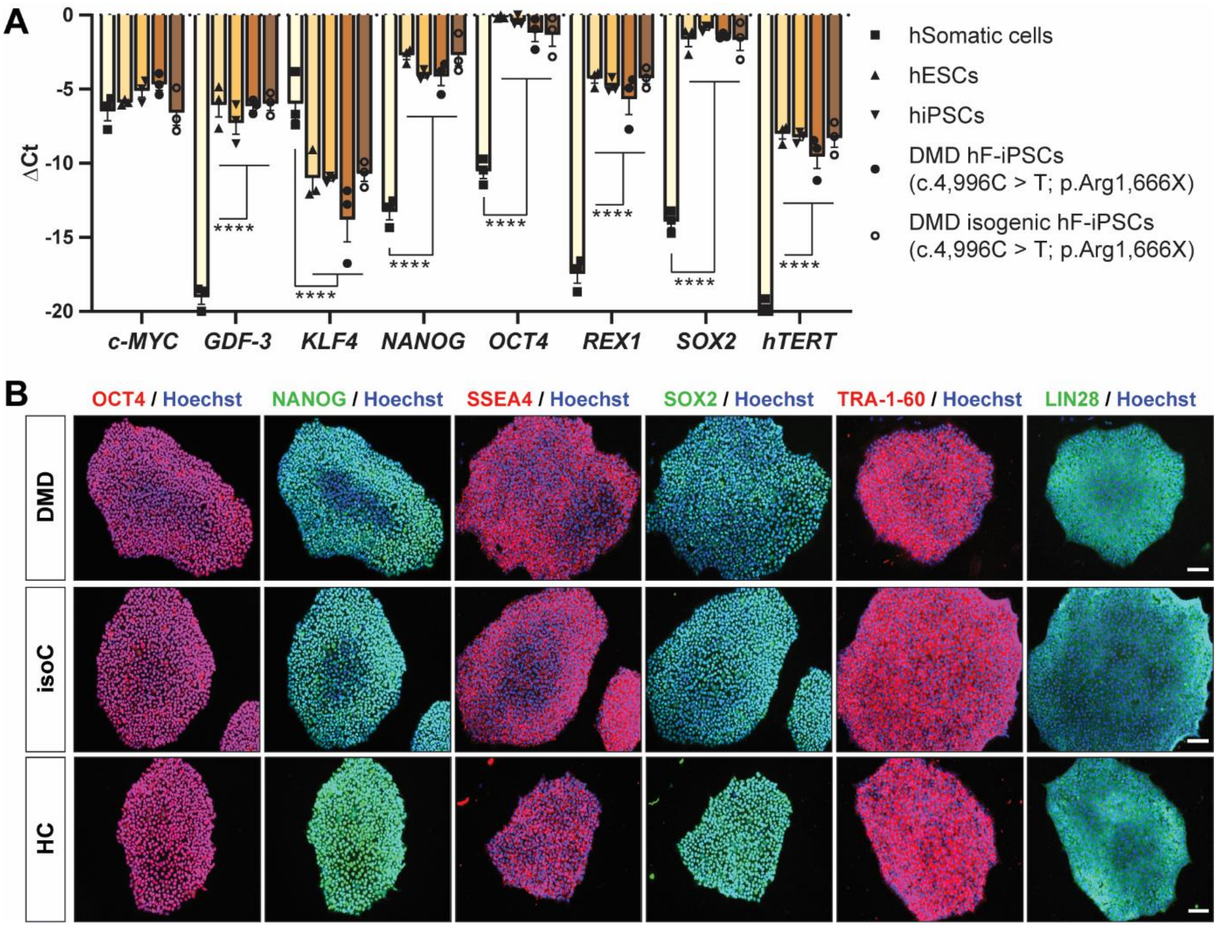


**Fig. S2: Characterization of the pluripotency state of the CRISPR/Cas9 corrected DMD isogenic control line.** The pluripotency state of the DMD isogenic control line. The following pluripotency genes (*c-MYC*, *GDF-3*, *KLF4*, *NANOG*, *OCT4*, *REX1*, *SOX2* and *hTERT*) **(A)** and proteins (OCT4, NANOG, SSEA4, SOX2, TRA-1-60 and LIN28) **(B)** were analyzed. Human embryonic stem cell lines (ESCs) and commercially available undifferentiated human iPSC lines were used as positive controls. Each data point was represented as ΔCt, normalized for the housekeeping genes (*GAPDH*, *HPRT* and *RPL13a*). Data were representative of three independent experiments (N = 3) and values were expressed as mean ± SEM. Significance of the difference was indicated as follows: *P < 0.05; **P < 0.01; ***P < 0.001 and ****P < 0.0001. Scale bar = 100 μm.

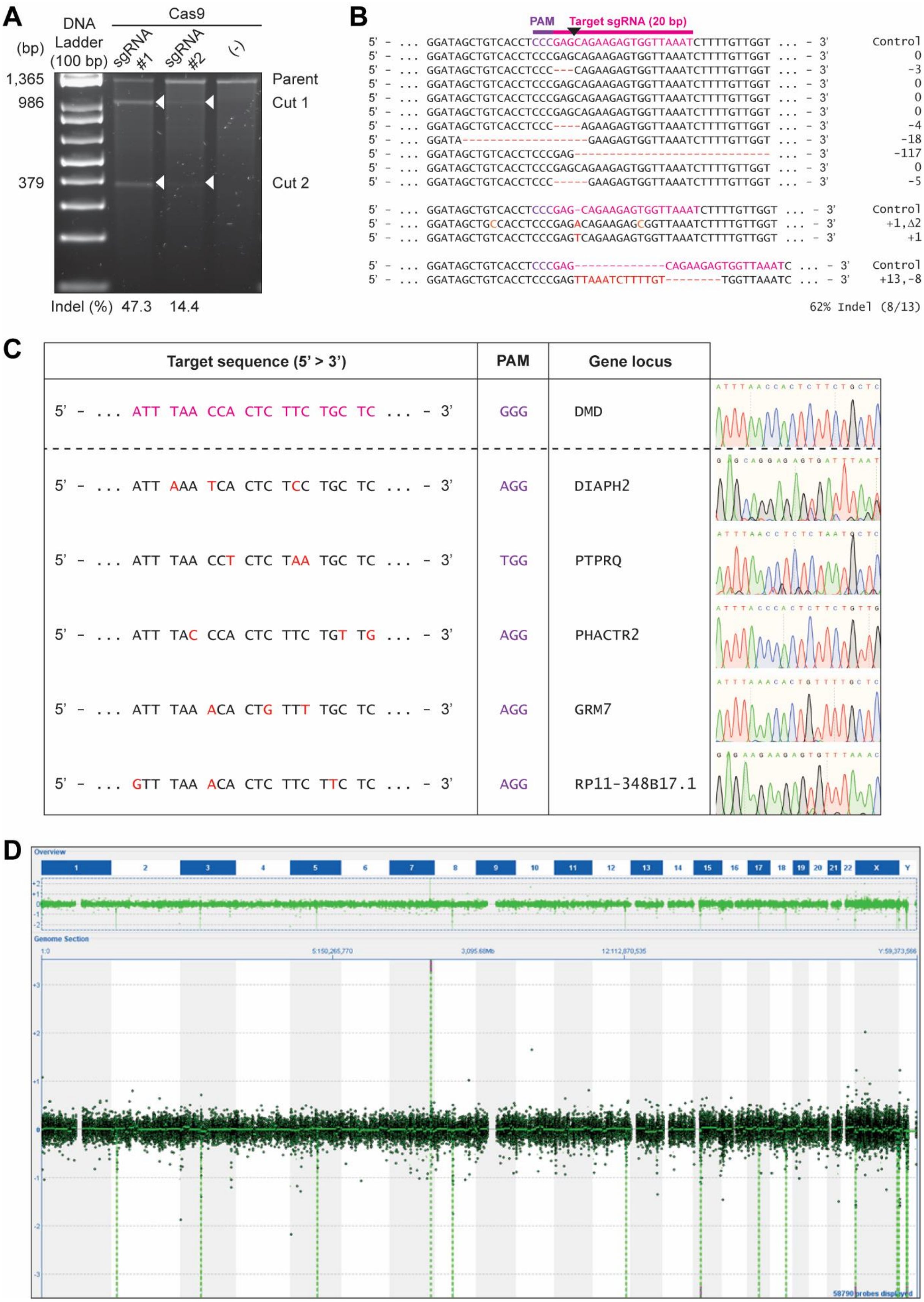


**Fig. S3: Validation of the Cas9 cutting efficiency and analysis of the off-targets. (A)** Surveyor assay in HEK293T cells to evaluate the cutting efficiency of the sgRNAs, represented as random events of base pair (bp) insertions or deletions (indel) after DSB. **(B)** DNA sequencing of the NHEJ events after transfection of the sgRNA-Cas9 plasmids in HEK293T cells. **(C)** List of CRISPR/Cas9 off-targets (source: www.synthego.com). **(D)** Detailed CGH molecular karyotyping showing no additional chromosomal abnormalities due to unwanted Cas9-mediated DSB cuts.

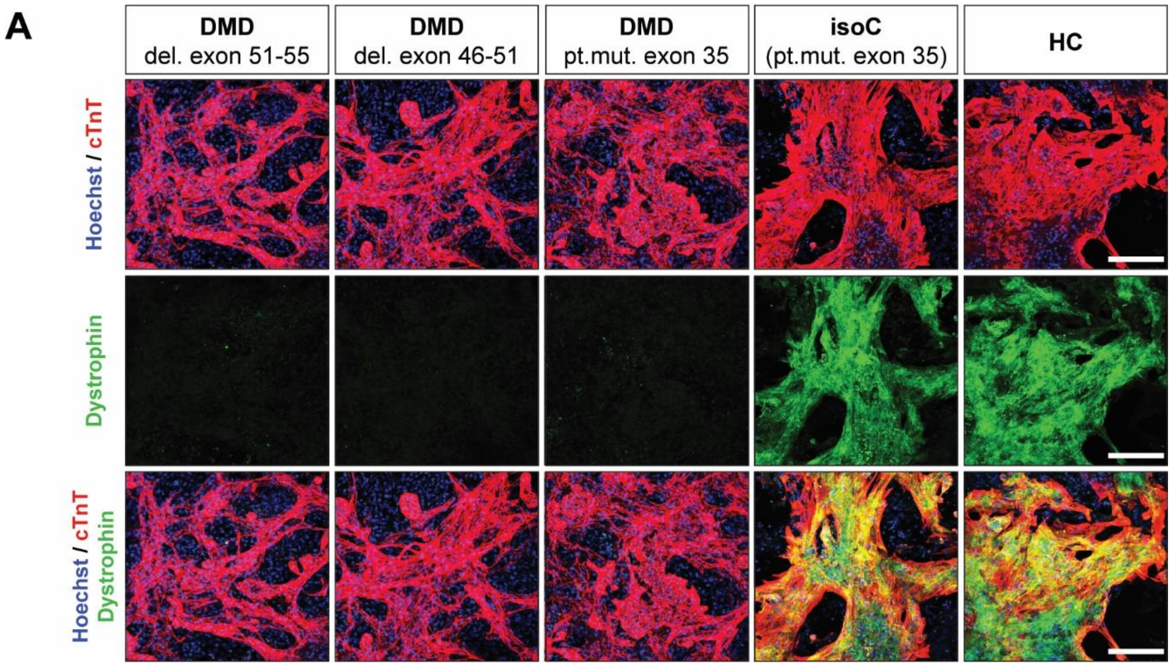


**Fig. S4: Lack of Dystrophin protein in iPSC-CMs from DMD patients resulted in premature cell death. (A)** Immunostaining showing the Dystrophin protein expression levels (green) in cTnT positive iPSC-CMs (cTnT, red and Hoechst, blue), derived from three DMD patient subjects (DMD #2: pt. mut. exon 35; DMD #5: del. exon 51-55 and DMD #6: del. exon 46-51) and controls (see also Table 1). Scale bar = 100 μm.

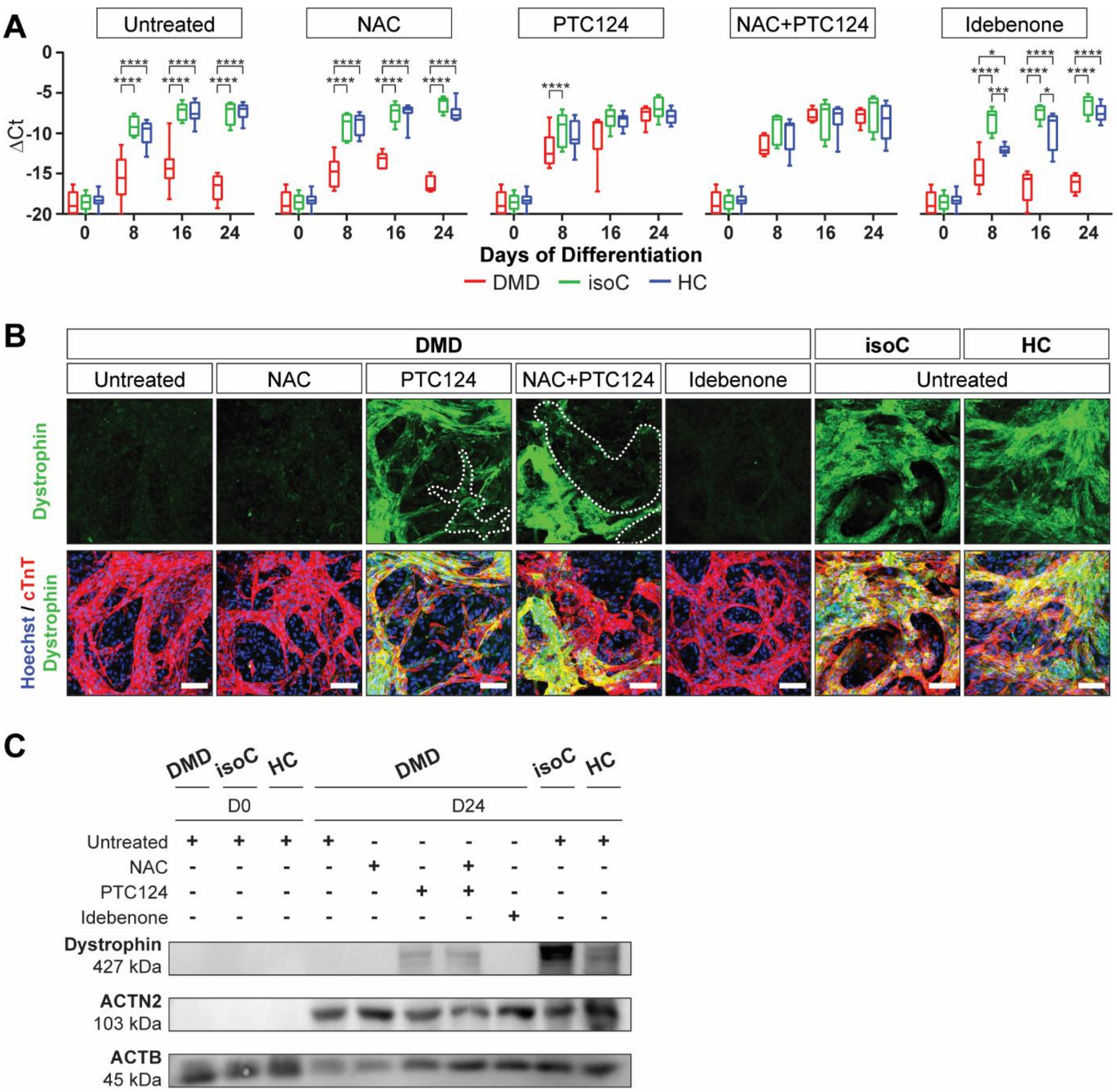


**Fig. S5: Dystrophin re-expression in DMD iPSC-CMs after PTC124 treatment. (A)** *Dystrophin* gene expression profiles in DMD iPSC-CMs, characterized by a genetic point mutation in exon 35 (c.4,996C > T; p.Arg1,666X) of the *Dystrophin* gene, upon NAC, PTC124 and idebenone addition. Each data point was represented as ΔCt, normalized for the housekeeping genes (*GAPDH* and *RPL13a*). Data were representative of five or more independent experiments (N ≥ 5) and values were expressed as mean ± SEM. Significance of the difference was indicated as follows: *P < 0.05; **P < 0.01; ***P < 0.001 and ****P < 0.0001 vs. subjects within the treatment condition. **(B)** Immunostaining at day 24 of differentiation demonstrating Dystrophin protein re-expression (green) upon PTC124 treatment in cTnT positive DMD and control iPSC-CMs (cTnT, red and Hoechst, blue). Scale bar = 100 μm. **(C)** Western blot analysis quantifying Dystrophin proteins in ACTN2 positive DMD and control iPSC-CMs, normalized to the loading protein ACTB.

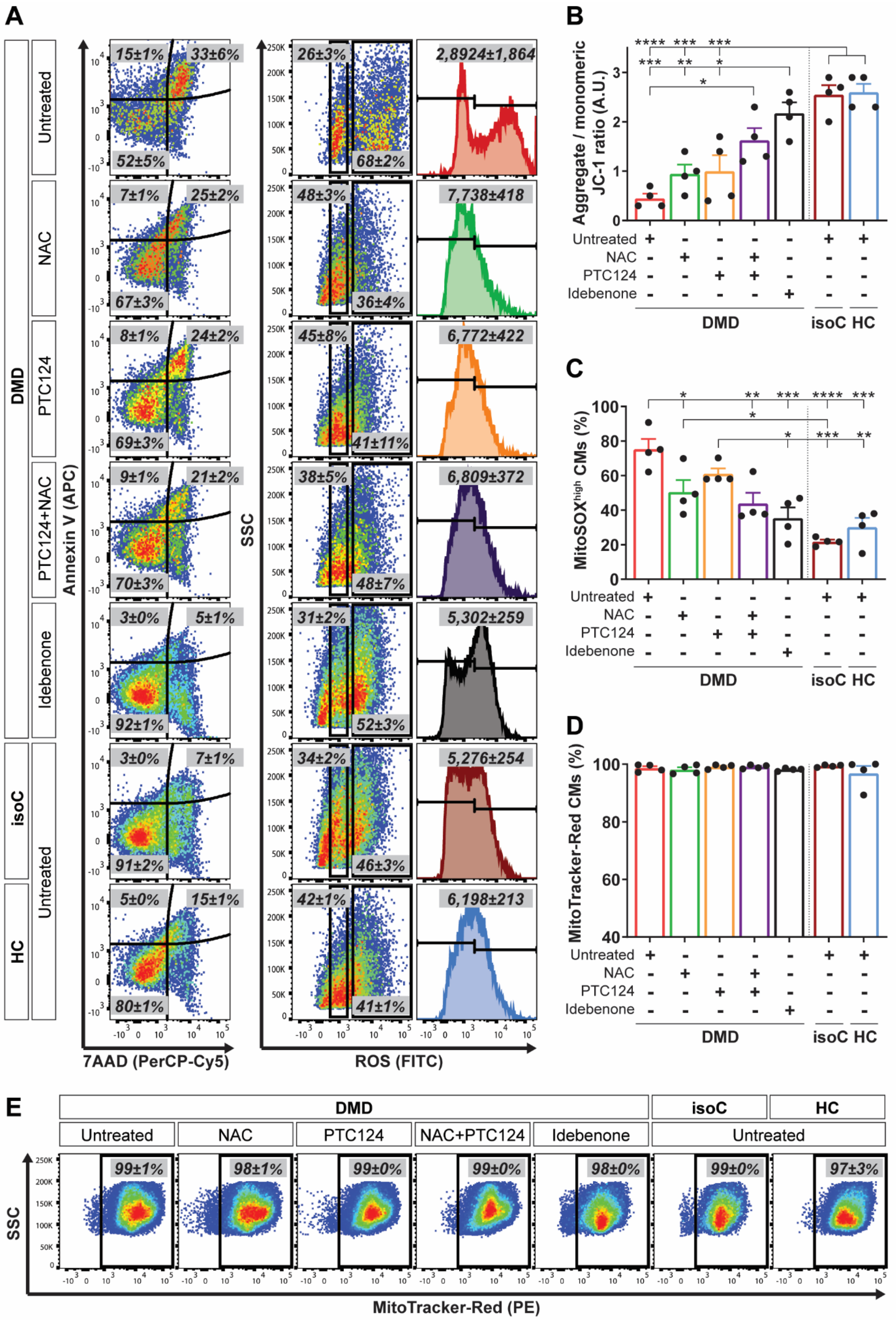
A
B
C
D
E
DMD
isoC
HC
Untreated
NAC
PTC124
PTC124+NAC
Idebenone
Annexin V (APC)
7AAD (PerCP-Cy5)
SSC
ROS (FITC)
Aggregate / monomeric JC-1 ratio (A.U.)
MitoSOX^high CMs (%)
MitoTracker-Red CMs (%)
MitoTracker-Red (PE)
NAC+PTC124
isoC HC

103

**Fig. S6: Corresponding flow cytometric graphs and quantification for the characterization of the cardiomyopathic phenotype of DMD iPSC-CMs, showing premature cell death, depolarized mitochondria and increased intracellular ROS levels.** **(A)** Representative flow cytometric analyses at day 15 of cardiac differentiation showing the percentage of cell death (using annexin V, APC and 7AAD, PerCP-Cy5, *left panels*) and intracellular ROS (FITC, *right panels*) in untreated and treated DMD iPSC-CMs compared to the DMD isogenic and healthy controls. iPSC-CMs were stained for SIRPA (PE) to obtain high CM purity (data not shown). Flow cytometric quantification at day 15 of differentiation showing the JC-1 aggregates/monomers ratio **(B)** and the mitochondrial superoxide production (MitoSOX) in depolarized DMD mitochondria **(C)** compared to DMD isogenic and healthy controls. **(D)** Percentage of MitoTracker-Red positive CMs upon NAC, PTC124 and idebenone treatment. **(E)** Corresponding flow cytometric analyses of the percentage of MitoTracker-Red (PE) positive iPSC-CMs. Data were representative of four independent experiments (N = 4). Data were reported as mean ± SEM. Significance of the difference was indicated as *P < 0.05; **P < 0.01; ***P < 0.001 and ****P < 0.0001.

119

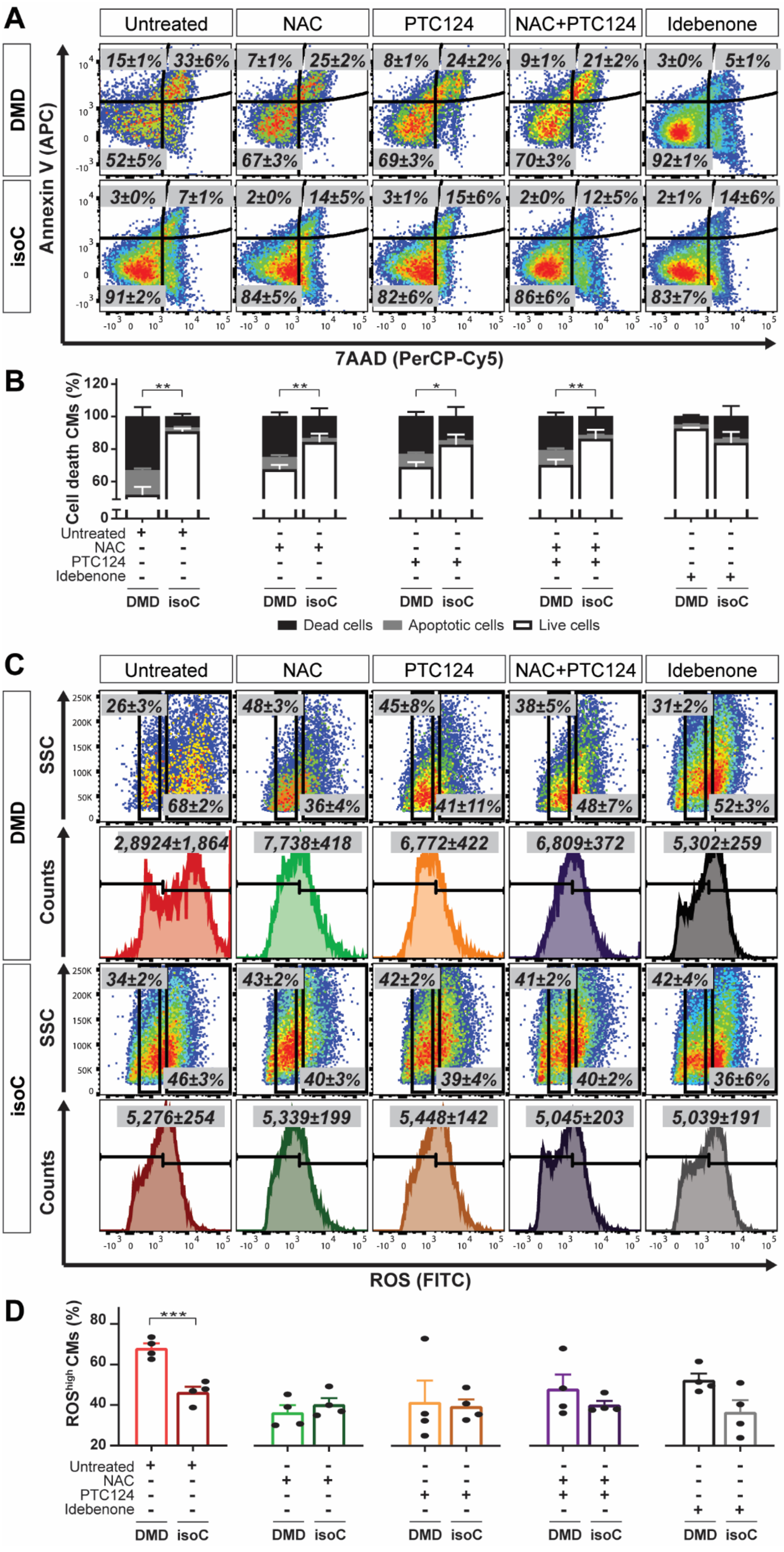

A
Untreated
NAC
PTC124
NAC+PTC124
Idebenone
DMD
isoC
Annexin V (APC)
7AAD (PerCP-Cy5)
B
Cell death CMs (%)
Untreated
NAC
PTC124
Idebenone
DMD
isoC
Dead cells
Apoptotic cells
Live cells
C
SSC
Counts
ROS (FITC)
D
ROS^high CMs (%)

**Fig. S7: Drug specificity on cell death and intracellular ROS concentrations in the experimental iPSC-CM groups. (A)** Example of flow cytometric analysis at day 15 of cardiac differentiation showing the percentage of cell death (using annexin V, APC and 7AAD, PerCP-Cy5) upon treatment in SIRPA (PE) positive iPSC-CMs derived from DMD and DMD isogenic controls. **(B)** Corresponding flow cytometric quantification for cell death observed after the treatment options. **(C)** Representative flow cytometric analyses showing intracellular ROS concentrations in DMD and DMD isogenic iPSC-CMs. **(D)** Quantification of the corresponding flow cytometric analyses showing the intracellular ROS levels. Data were representative of four independent experiments (N = 4). Data were reported as mean ± SEM. Significance of the difference was indicated as *P < 0.05; **P < 0.01; ***P < 0.001 and ****P < 0.0001.

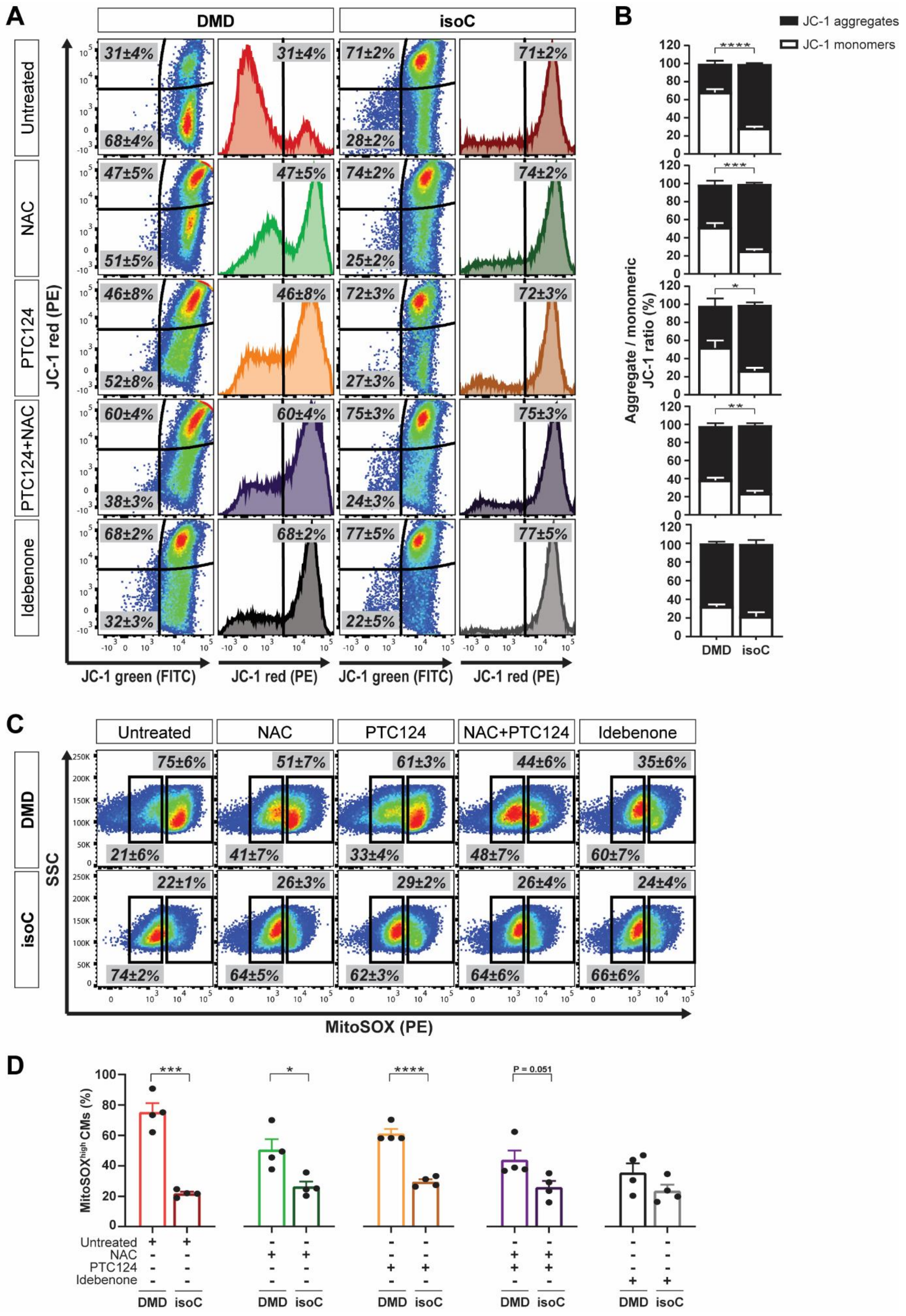
A
DMD
isoC
Untreated
NAC
PTC124
PTC124+NAC
Idebenone
JC-1 red (PE)
JC-1 green (FITC)
31±4%
68±4%
47±5%
51±5%
46±8%
52±8%
60±4%
38±3%
68±2%
32±3%
71±2%
28±2%
74±2%
25±2%
72±3%
27±3%
75±3%
24±3%
77±5%
22±5%
B
JC-1 aggregates
JC-1 monomers
Aggregate / monomeric JC-1 ratio (%)
DMD
isoC
C
Untreated
NAC
PTC124
NAC+PTC124
Idebenone
DMD
isoC
SSC
MitoSOX (PE)
75±6%
21±6%
51±7%
41±7%
61±3%
33±4%
44±6%
48±7%
35±6%
60±7%
22±1%
74±2%
26±3%
64±5%
29±2%
62±3%
26±4%
64±6%
24±4%
66±6%
D
MitoSOX$^{high}$ CMs (%)
P = 0.051
Untreated
NAC
PTC124
Idebenone
DMD
isoC

**Fig. S8: Drug specificity on $\Delta\Psi_m$ and mitochondrial superoxide concentrations in the experimental iPSC-CM groups. (A)** Representative flow cytometric analyses at day 15 of cardiac differentiation for JC-1 aggregates (PE) and JC-1 monomers (FITC) upon treatment in DMD iPSC-CMs and the DMD isogenic counterpart. **(B)** Corresponding flow cytometric quantification for $\Delta\Psi_m$. **(C)** Flow cytometric analyses at day 15 of differentiation showing the mitochondrial superoxide production (MitoSOX, PE) in depolarized DMD mitochondria. **(D)** Quantification of the corresponding flow cytometric analyses showing the mitochondrial superoxide production (MitoSOX). Data were representative of four independent experiments (N = 4). Flow cytometry data were reported as mean ± SEM. Significance of the difference was indicated as follows: *P < 0.05; **P < 0.01; ***P < 0.001 and ****P < 0.0001.

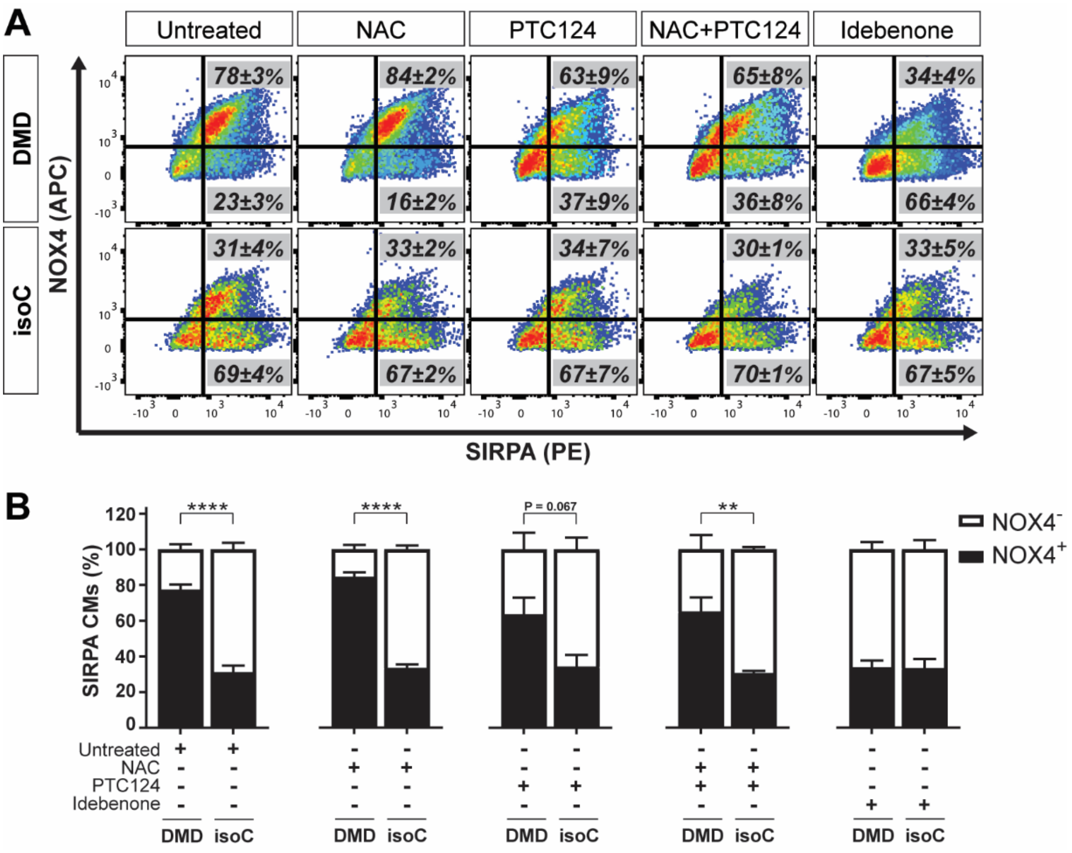


**Fig. S9: The specificity of the treatment options on the expression levels of NOX4 in iPSC-CM cultures. (A)** Example of flow cytometric analysis at day 15 of cardiac differentiation showing the percentage of NOX4 (APC) on SIRPA (PE) positive DMD and DMD isogenic iPSC-CMs upon the treatment options. **(B)** Corresponding flow cytometric quantification for NOX4. Data were representative of three independent experiments (N = 3). Flow cytometry data were reported as mean ± SEM. Significance of the difference was indicated as follows: *P < 0.05; **P < 0.01; ***P < 0.001 and ****P < 0.0001.

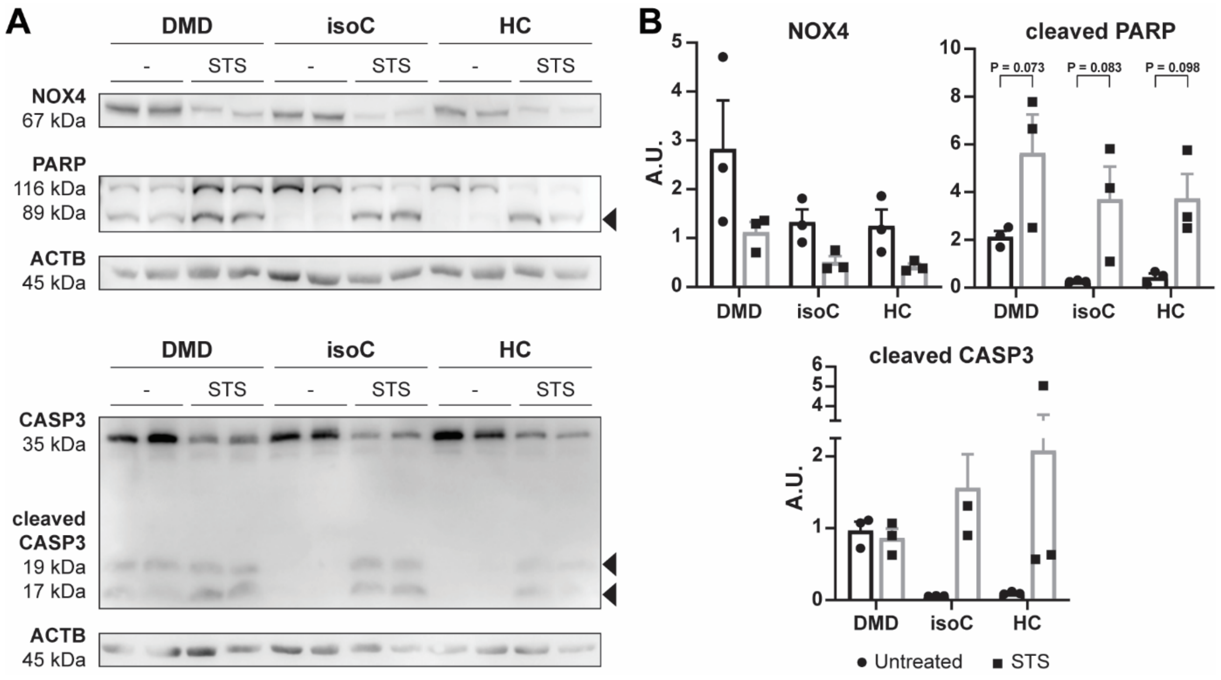


**Fig. S10: NOX4 protein expression levels after STS-induced cell death in DMD and control iPSC-CMs. (A)** Western blot analysis showing cardiac NOX4 proteins and the cell death markers Poly (ADP-ribose) polymerase (PARP) and Caspase-3 (CASP3) in DMD and control iPSC-CMs after a 6 h exposure to 1 μM STS. Cleaved forms of PARP and CASP3 are indicated by black triangles. ACTB was used as loading control. **(B)** Quantification of the western blot analysis for the markers NOX4, cleaved PARP and cleaved CASP3. Data were representative of three independent experiments (N = 3) and values were expressed as mean ± SEM.

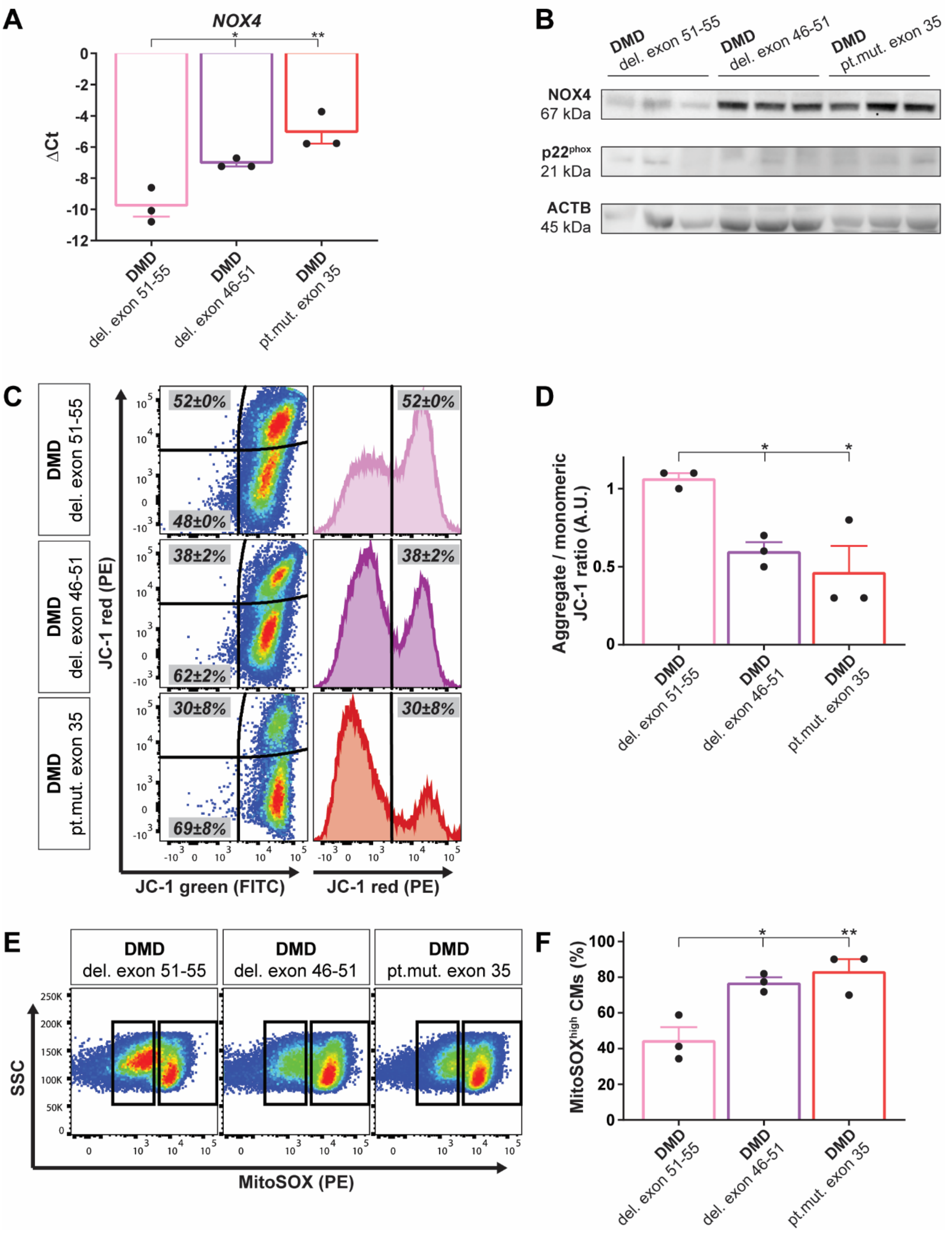


**Fig. S11: Characterization of DMD patient-specific iPSC-CMs *in vitro*, showing increased NOX4 gene and protein expression levels, depolarized mitochondria and increased intracellular ROS levels. (A)** *NOX4* mRNA levels in iPSC-CMs of three DMD patients at day

8 of cardiac differentiation. Each data point was represented as ΔCt, normalized for the housekeeping genes (*GAPDH* and *RPL13a*). Data were representative of three independent experiments (N = 3) and values were expressed as mean ± SEM. Significance of the difference was indicated as follows: *P < 0.05; **P < 0.01; ***P < 0.001 and ****P < 0.0001. **(B)** Western blot analysis quantifying the corresponding NOX4 protein levels and its regulatory subunit $p22^{phox}$, normalized to the loading protein ACTB. **(C)** Representative flow cytometric analyses at day 15 of differentiation for JC-1 aggregates (PE) and JC-1 monomers (FITC) in three DMD patient-specific iPSC-CMs. **(D)** Corresponding flow cytometric quantification of JC-1 aggregates and JC-1 monomers. **(E)** Flow cytometric analyses at day 15 of differentiation showing the mitochondrial superoxide production (MitoSOX, PE) in depolarized DMD mitochondria. **(F)** Corresponding flow cytometric quantification for the number of CMs with high mitochondrial superoxide concentrations. Data were representative of three independent experiments (N = 3). Flow cytometry data were reported as mean ± SEM. Significance of the difference was indicated as follows: *P < 0.05; **P < 0.01; ***P < 0.001 and ****P < 0.0001.

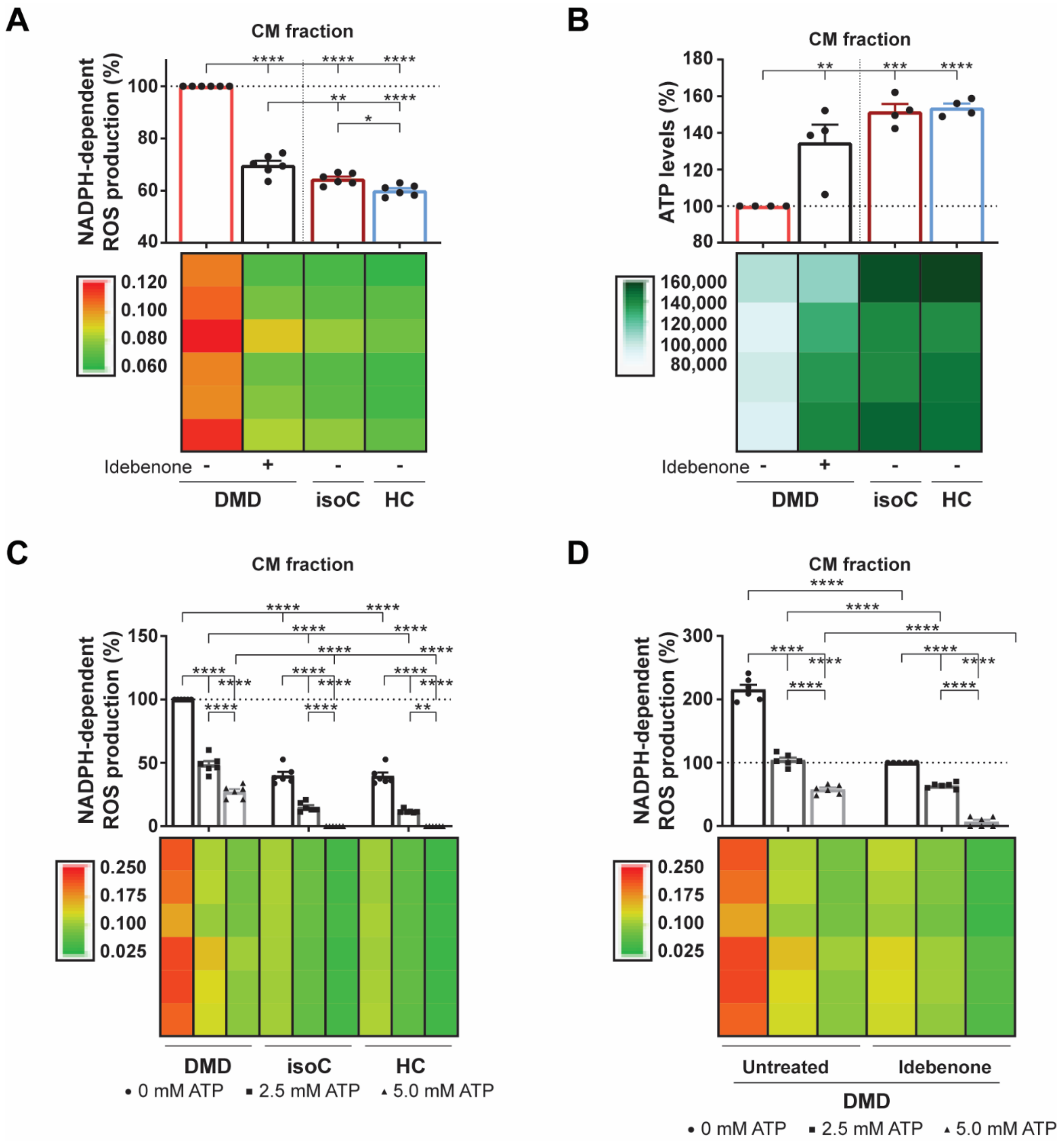


**Fig. S12: NADPH-dependent ROS production and intracellular ATP levels in DMD iPSC-CMs after idebenone application. (A)** Quantification of the NADPH-dependent superoxide production of NOX4 in the total CM fraction of DMD iPSC-CMs with or without idebenone addition compared to controls. **(B)** ATP luminescence detection showing the effect of idebenone treatment on the intracellular ATP levels in DMD iPSC-CMs. **(C)** Quantification of the ROS-producing NOX4 activity after 2.5 and 5.0 mM ATP addition in DMD iPSC-CM and control cultures. Each data point was represented as percentage (%), normalized to the total CM fraction of the untreated DMD iPSC-CMs. **(D)** Quantification of the NADPH-dependent superoxide production of NOX4 in the total CM fraction of DMD iPSC-CMs upon 2.5 and 5.0 mM ATP addition, with or without idebenone treatment. Each data point was represented as percentage (%), normalized to the total CM fraction of the idebenone-treated DMD iPSC-CM cultures. Data were representative of four or six independent experiments (N = 4 or N = 6) and values were expressed as mean ± SEM. Colored rectangles represented the independent experiments. Significance of the difference was indicated as follows: *P < 0.05; **P < 0.01; ***P < 0.001 and ****P < 0.0001.

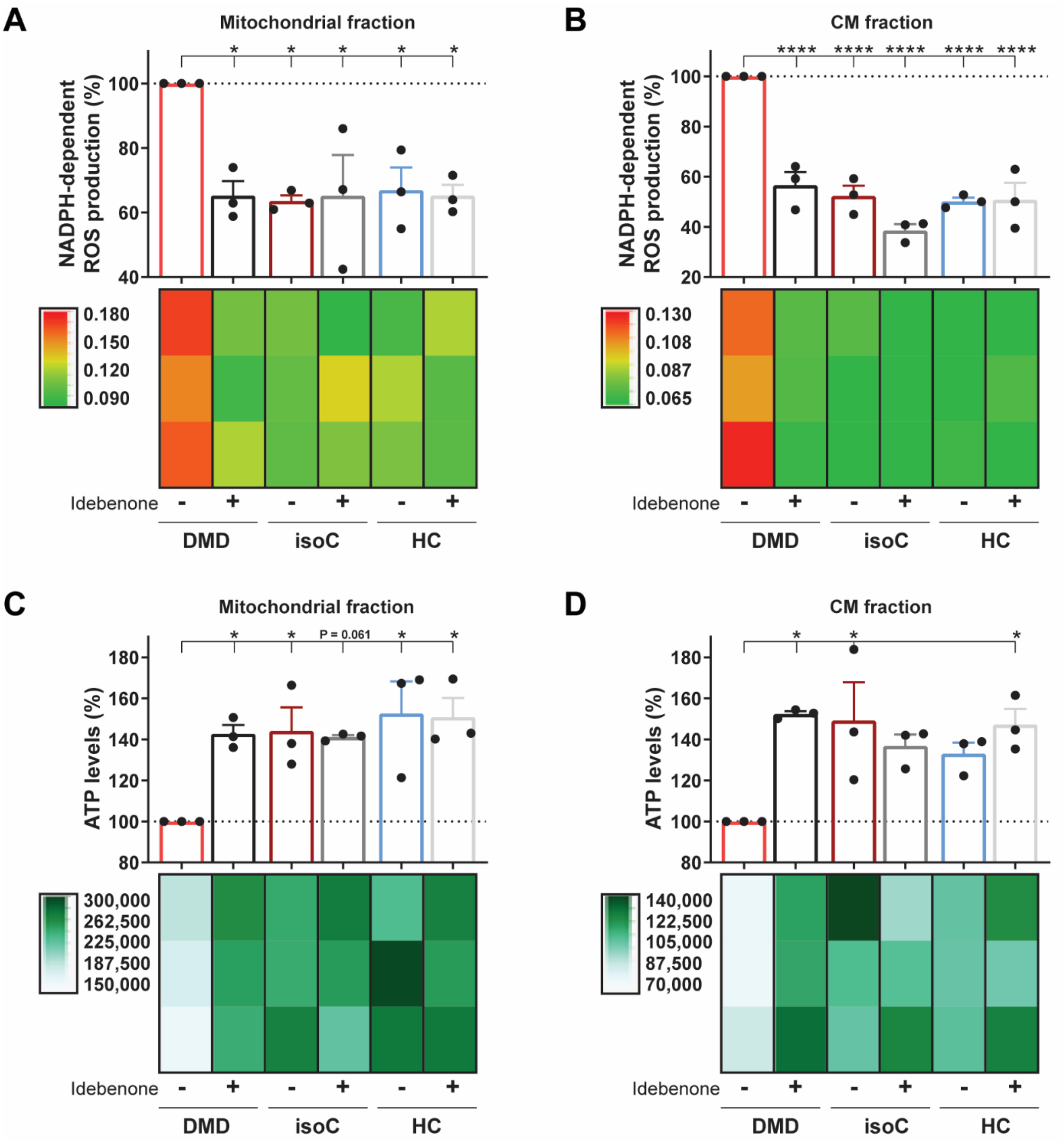


**Fig. S13: The specificity of idebenone on the NADPH-dependent ROS production and ATP levels in the experimental iPSC-CM groups.** Quantification of the NADPH-dependent superoxide production of NOX4 in the mitochondrial **(A)** and CM fraction **(B)** of iPSC-CMs derived from DMD, DMD isogenic and healthy controls with or without idebenone treatment. **(C-D)** ATP luminescence detection showing the effect of idebenone treatment on the ATP levels in iPSC-CMs cultures. Each data point was represented as percentage (%), normalized to the untreated DMD iPSC-CM cultures. Data were representative of three independent experiments (N = 3) and values were expressed as mean ± SEM. Colored rectangles represented the independent experiments. Significance of the difference was indicated as follows: *P < 0.05; **P < 0.01; ***P < 0.001 and ****P < 0.0001.
